# Evolution and instability of human centromeres are accelerated by heterochromatin boundary loss and CENP-A overexpression

**DOI:** 10.1101/2025.02.03.636285

**Authors:** Megan A. Mahlke, Lior Lumerman, Poulomi Nath, Cy Chittenden, Savannah Hoyt, Jonas Koeppel, Yuan Xu, Rebecca Raphael, Kylie Zaffina, Paul W. Hook, Winston Timp, Karen H. Miga, Peter J. Campbell, Rachel J. O’Neill, Nicolas Altemose, Yael Nechemia-Arbely

## Abstract

Centromere location is specified by CENP-A, a centromere-specific histone that epigenetically defines centromere identity. How CENP-A is maintained at one location in rapidly evolving centromeric DNA is unknown. Using single-cell-derived clones of human cell lines, we demonstrate single-cell heterogeneity in CENP-A position within cell populations at neocentromeres and a native centromere. CENP-A heterogeneity is accompanied by unique DNA methylation and H3K9me3 patterns, with DNA methylation shifting according to CENP-A position. We further demonstrate centromere epigenetic evolution over prolonged proliferation, with native centromeres maintaining stable heterochromatin boundaries, but neocentromeres exhibiting DNA methylation instability, H3K9me3 gain, boundary loss and fragility. Lastly, prolonged CENP-A and HJURP overexpression leads to centromere and neocentromere expansion, gradual CENP-A depletion, neocentromere destabilization and CENP-A re-localization that is accompanied by local heterochromatin remodeling. This study reveals the naturally evolving epigenetic plasticity of human centromeres and neocentromeres and highlights the importance of repressive chromatin boundaries in maintaining centromere stability.

## Introduction

Centromeres play a crucial role in preserving genomic stability by ensuring that chromosomes are properly segregated into daughter cells. In humans, centromeres complete this formidable task through precise epigenetic maintenance of histone H3 variant CENP-A that defines centromere identity at one location per chromosome (Jansen et al., 2007; Mahlke and Nechemia-Arbely, 2020; Nechemia-Arbely et al., 2019; Sullivan et al., 1994; Warburton et al., 1997). CENP-A recruits CENP-C that then nucleates the constitutive centromere associated network (CCAN) (Foltz et al., 2006), ultimately leading to kinetochore formation and proper sister chromatid segregation. Disrupting CENP-A regulation through its overexpression or knockdown can cause ectopic CENP-A accumulation and/or weakening of endogenous centromeres, resulting in chromosomal aneuploidies and genomic instability (Black et al., 2007; Fachinetti et al., 2013; Lacoste et al., 2014; Li and Zhu, 2022; Mahlke and Nechemia-Arbely, 2020; Nechemia-Arbely et al., 2019; Shrestha et al., 2017; Shrestha et al., 2021). Therefore, the deposition and maintenance of CENP-A are tightly regulated. CENP-A deposition at centromeres by its chaperone HJURP occurs once per cell cycle during early G1 (Dunleavy et al., 2009; Foltz et al., 2009; Jansen et al., 2007; Nechemia-Arbely et al., 2012) and is both negatively regulated by CDKs and promoted by PLK1 (Conti et al., 2024) providing strict temporal control. During DNA replication, errant ectopically deposited CENP-A is stripped from chromosome arms but centromeric CENP-A is selectively maintained by the coordinated action of MCM2 helicase, CAF1, HJURP, and the CCAN (Nechemia-Arbely et al., 2019; Zasadzińska et al.), enabling precise CENP-A reassembly onto both daughter centromeres (Mahlke et al., 2023; Nechemia-Arbely et al., 2019).

Human centromeres are found at tandemly repeating α-satellite High Order Repeat (HOR) arrays (Altemose et al., 2022a; Logsdon et al., 2024). CENP-A is localized to a specific position on each active HOR array per chromosome and is maintained at the same centromeric sequences throughout the cell cycle with precision (Mahlke et al., 2023; Nechemia-Arbely et al., 2019; Ross et al., 2016). However, the idea of precise CENP-A maintenance is at odds with the highly evolving nature of centromeric HOR α-satellite arrays (Altemose et al., 2022a; Logsdon et al., 2024) and the discovery of differential CENP-A placement at the same centromeres in human populations (Mahlke et al., 2023; Volpe et al., 2023). CENP-A position at non-repetitive centromeres can drift between maize species, yeast strains, horse generations, and within a single population of chicken cells (Allshire et al., 1994; Gent et al., 2017; Hori et al., 2016; Purgato et al., 2015). Humans can acquire non-repetitive centromeres called neocentromeres through relocation of CENP-A to chromosome arms (Amor and Choo, 2002; Fukagawa and Earnshaw, 2014; Warburton, 2004), but to what extent CENP-A position drifts at established human neocentromeres or native repetitive centromeres is unknown.

Human centromeres are characterized by a unique epigenetic environment, wherein α-satellite HORs are highly methylated but feature a decrease in DNA methylation at the centromere dip region (CDR) where CENP-A is enriched and the kinetochore assembles (Altemose et al., 2022a; Gershman et al., 2022; Logsdon et al., 2021; Miga et al., 2020). Human centromeres are also flanked by pericentromeric heterochromatin enriched for H3K9me3 (Gershman et al., 2022; McKinley and Cheeseman, 2016; Sullivan and Karpen, 2004) that can suppress CENP-A assembly (Ohzeki et al., 2016; Ohzeki et al., 2012) and is thought to function as a centromere boundary (Lam et al., 2006; McKinley and Cheeseman, 2016; Ohzeki et al., 2016; Ohzeki et al., 2012) that stabilizes CENP-A position (Martins et al., 2016; Murillo-Pineda et al., 2021; Scott et al., 2006; Slee et al., 2012). At human neocentromeres, H3K9me3 can contribute to establishing heterochromatin boundaries (Lam et al., 2006; Murillo-Pineda et al., 2021; Naughton et al., 2022). Whether DNA methylation dips are present at neocentromeres and their possible contribution to neocentromere stability remains unknown.

The paradox between precise CENP-A maintenance and diverse, rapidly evolving human centromere sequences led us to investigate whether CENP-A position is precisely maintained at centromeres of single cells within a cell population over prolonged proliferation. Using human cell lines that harbor non-repetitive fully functional neocentromeres, we reveal that CENP-A position is heterogenous at neocentromeres within a population of cells and these unique CENP-A positions are precisely maintained for at least ∼40 cell cycles. Over prolonged proliferation (∼280 cell cycles), unique CENP-A positions evolve, generating new cell populations with increased CENP-A positional heterogeneity. Using Directed Methylation with Long-read sequencing (DiMeLo-seq) (Altemose et al., 2022b), we also find single-cell clonal CENP-A heterogeneity at a native centromere. CENP-A heterogeneity is accompanied by heterogenous DNA methylation and H3K9me3 patterns at centromeres and neocentromeres. These findings are paradigm shifting, as CENP-A has been observed as a stable, long-lived nucleosome that is inherited indefinitely and precisely to maintain and propagate centromere identity (Fachinetti et al., 2013; Mahlke et al., 2023; Nechemia-Arbely et al., 2019; Ross et al., 2016; Smoak et al., 2016).

We are the first to report a DNA methylation CDR at a human neocentromere. We further show that the CDR of a centromere or neocentromere can vary in single cells in accordance with CENP-A position. Over prolonged proliferation, native centromeres maintain a DNA methylation CDR at the site of CENP-A enrichment but neocentromeres fail to maintain their CDRs and gain H3K9me3 at sites of CENP-A enrichment, leading to boundary loss, increased epigenetic heterogeneity and centromere fragility. Finally, we demonstrate that low level CENP-A and HJURP overexpression causes centromere and neocentromere expansion but prolonged elevated overexpression of CENP-A and HJURP is sufficient to destabilize an existing neocentromere and its CDR, prompting the formation of a new neocentromere at a different chromosome locus through heterochromatin remodeling. This work highlights the role of the DNA methylation CDR in confining the evolution of centromere epigenetic identity and thereby maintaining the stability of human centromeres.

## Results

### CENP-A position at neocentromeres is heterogenous within a cell population

To study stability of CENP-A position at human centromeres, we used the PD-NC4 cell line that harbors a stable neocentromere (NeoCEN4) at a non-repetitive region of the q arm of one copy of chromosome 4, enabling high-resolution mapping of CENP-A (Amor et al., 2004) (Fig. 1A). NeoCEN4 is located in band 4q22.1 and has three distinct CENP-A peaks spanning a ∼250 kb region (Hasson et al., 2013). We generated 9 single-cell derived PD-NC4 clones and confirmed that clones retained NeoCEN4 (Fig. S1A, Table S1). CENP-A ChIP-seq confirmed three CENP-A peaks at NeoCEN4 in PD-NC4 parental cells (Fig. 1A, B). However, single-cell derived PD-NC4 clones exhibited CENP-A positions that differed from the expected three-peak pattern (Fig. 1B) and were classified into three distinct CENP-A positional phenotypes: maintaining all three CENP-A peaks (similar to parental population, Fig. 1B, blue), loss of CENP-A peak 1 (Fig. 1B, green) or loss of CENP-A peak 3 (Fig. 1B, red). Whole genome sequencing for three clones best representing the three CENP-A position phenotypes (clones C2, D2 and E2) confirmed that distinct CENP-A positions were true epigenetic events and not a result of genomic rearrangements at NeoCEN4 (Fig. S1D). Identifying regions significantly enriched for CENP-A (p<0.00001, fold change (fc) >20) in parental PD-NC4 cells and in clones shows the position of CENP-A peaks in clones can significantly drift (Fig. 1B, clone C4, Peak 3), or expand (Fig. 1B, clones D1, A1, C4, E1, E2) in the range of 12 – 25 kb per CENP-A peak beyond CENP-A position observed in the parental PD-NC4 population (Fig. 1B). Clones also exhibited mild CENP-A peak shrinkage of 3 – 5 kb (Clones C2, C3, D1) and convergence of remaining CENP-A peaks (Clones C4, E1 and E2). Thus, the position of CENP-A at NeoCEN4 observed in PD-NC4 parental cells represents a population average of distinct CENP-A positions.

**Figure 1.**
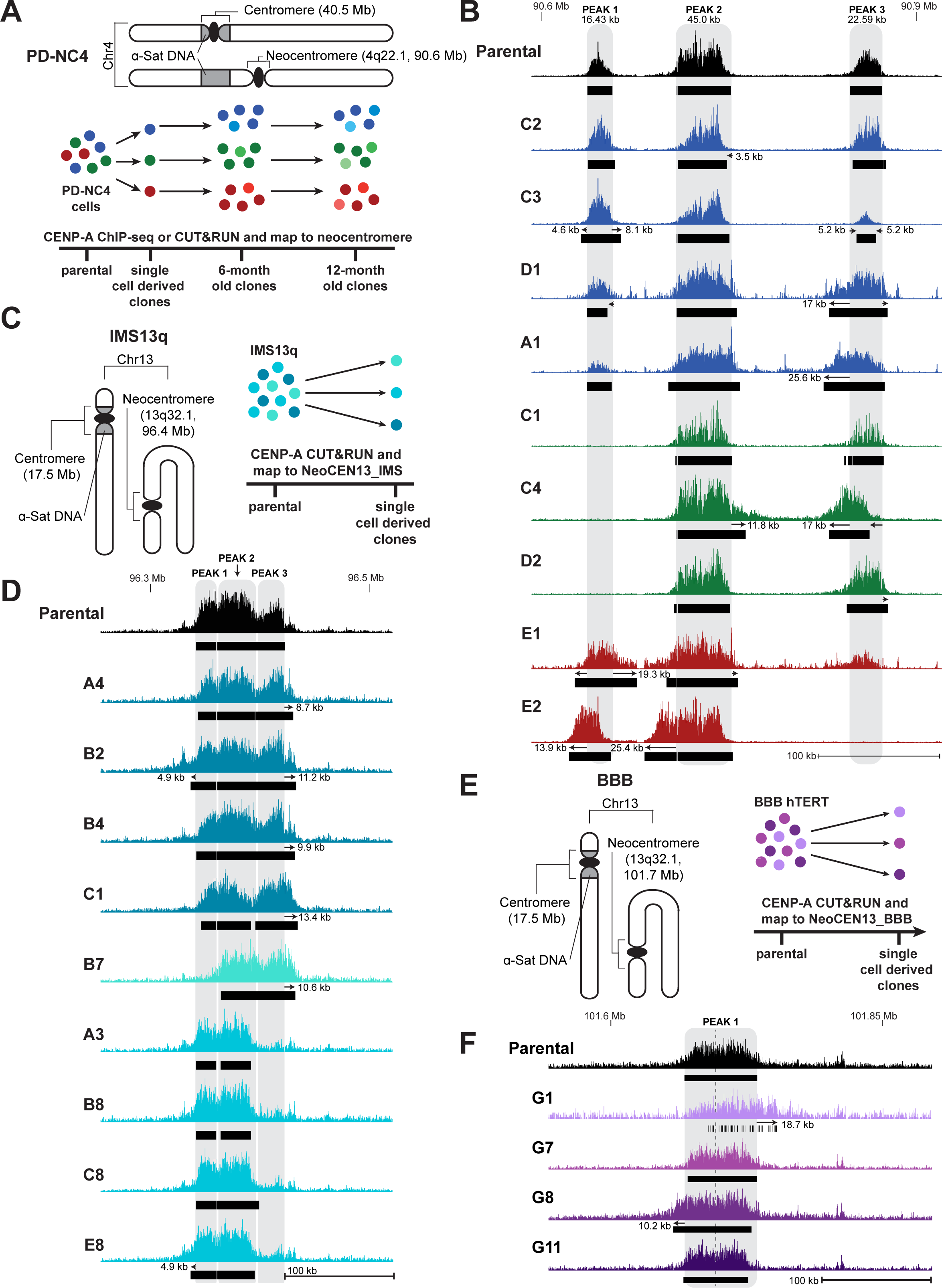
CENP-A position at neocentromeres is heterogenous within a population of cells. **A.** Diagram of both copies of chromosome 4 in PD-NC4 cells and experimental design. In the normal copy, CENP-A is located in the repetitive α-satellite DNA at 40.5 Mb (T2T CHM13v2). In the other copy, CENP-A has relocated to a non-repetitive site at 4q22.1, 90.6 Mb (T2T CHM13v2) and formed a neocentromere. Single-cell derived PD-NC4 clones were isolated and maintained in culture for 12 months. ChIP-seq or CUT&RUN for CENP-A was performed on clones when newly derived (young) and after 6 and 12 months. **B.** CENP-A ChIP-seq or CUT&RUN at NeoCEN4 in PD-NC4 parental cells (top, black) and in PD-NC4 single-cell derived clones. Clones are colored according to CENP-A positional phenotype: three CENP-A peaks (blue), loss of CENP-A peak 1 (green), loss of CENP-A peak 3 (red). Numbers above parental peaks indicate peak size. **C.** Diagram of both copies of chromosome 13 in IMS13q cells and experimental design. In the normal copy, CENP-A is located in the repetitive α-satellite DNA at 17.5 Mb. In the other copy, a chromosome break led to a fusion of two Chr13 q arms which then formed a neocentromere by acquiring CENP-A at the non-repetitive locus 13q32.1, 96.4 Mb. **D.** CENP-A CUT&RUN at NeoCEN13 in IMS13q parental cells (top, black) and in newly isolated IMS13q single-cell derived clones. Clones are colored according to CENP-A positional phenotype: three CENP-A peaks (teal), loss of CENP-A peak 1 (turquoise), loss of CENP-A peak 2 (aqua). **E.** Diagram of both copies of chromosome 13 in BBB cells and experimental design. In the normal copy, CENP-A is located in the repetitive α-satellite DNA at 17.5 Mb. In the other copy, a chromosome break led to a fusion of two Chr13 q arms which then formed a neocentromere by acquiring CENP-A at the non-repetitive locus 13q32.1, 101.7 Mb. **F.** CENP-A CUT&RUN at NeoCEN13 in BBB parental cells (top, black) and in newly isolated single-cell derived BBB clones. Dashed line in peak 1 indicates probable division into two CENP-A peaks. For panels **B, D,** and **F,** black bars show location of significantly enriched CENP-A peaks (p<0.00001, fc ≥ 20). Grey shadows indicate the position of significantly enriched CENP-A peaks in parental cell populations. Arrows indicate CENP-A drift, expansion or shrinkage (defined by significantly enriched peaks) in clones relative to parental cells.

To confirm whether CENP-A heterogeneity is a general feature of human neocentromeres, we derived single-cell clones from two additional human cell lines with non-repetitive neocentromeres on chromosome 13, IMS13q and BBB (Alonso et al., 2010) (Fig. 1C, E). CENP-A CUT&RUN in IMS13q and BBB parental cells and single-cell derived clones (Fig. 1C, E, Fig. S1B-C, Tables S2, S3) revealed three distinct CENP-A peaks significantly enriched for CENP-A (p<0.00001, fc >20) at the IMS13q neocentromere (NeoCEN13_IMS) (Fig. 1D, Parental), and a single CENP-A peak (p<0.00001, fc >20) at the BBB neocentromere (NeoCEN13_BBB), although CENP-A distribution suggests two adjacent domains (Fig. 1F, Parental). CENP-A position at the NeoCEN13 of IMS13q and BBB clones differed significantly from that of respective parental populations (Fig. 1D, F). We identified CENP-A drift at NeoCEN13_IMS up to 13 kb (clone C1), CENP-A expansion up to 16 kb (clones A4, B2, B4), and loss of CENP-A peak 1 (clone B7) or peak 3 (clones A3, B8, C8, E8). We identified CENP-A drift at NeoCEN13_BBB of up to 18 kb (clone G1). Thus, heterogeneity of CENP-A position within cell populations is a general feature of human neocentromeres.

### Changes in CENP-A position do not affect centromere function

Centromere function is impaired when centromeric CENP-A is depleted (Black et al., 2007; Fachinetti et al., 2013). To determine if loss of a CENP-A peak or CENP-A drift at neocentromeres correlates with changes in local CENP-A abundance, we measured the length of DNA significantly enriched for CENP-A (p<0.00001, fc >20) at neocentromeres in parental PD-NC4, IMS13q, and BBB cells and their respective single-cell derived clones. Parental PD-NC4 cells have CENP-A enrichment along 84 kb of DNA at NeoCEN4 (Table S4). Despite CENP-A loss, drift and expansion, the total length of DNA enriched for CENP-A at NeoCEN4 in PD-NC4 clones remained similar to or greater than parental PD-NC4 cells, ranging between 74 – 107 kb (Table S4). Quantification of CENP-A intensity at NeoCEN4 and translation into number of CENP-A molecules (Bodor et al., 2014) revealed no significant difference in CENP-A intensity between parental PD-NC4 cells and clones (Fig. 2A, B). Instead, a similar number of CENP-A molecules is maintained at NeoCEN4 in all PD-NC4 clones irrespective of CENP-A position. NeoCEN13_IMS and NeoCEN13_BBB had a total of 80.37 kb and 66.86 kb of DNA significantly enriched for CENP-A in parental cells, respectively (Tables S5, S6). IMS13q clones lost variable lengths of DNA enriched for CENP-A, with loss of peak 3 being most detrimental and total length of CENP-A enrichment in clones ranging between 46 – 95 kb (Table S5). In contrast, the length of CENP-A enriched region remained similar between BBB parental cells and clones at NeoCEN13, ranging between 60 – 70 kb (Table S6). These findings suggest that different neocentromeres may have variable tolerance for CENP-A plasticity, and that the minimal CENP-A enriched DNA size required for functional neocentromere assembly is ∼46 kb.

**Figure 2.**
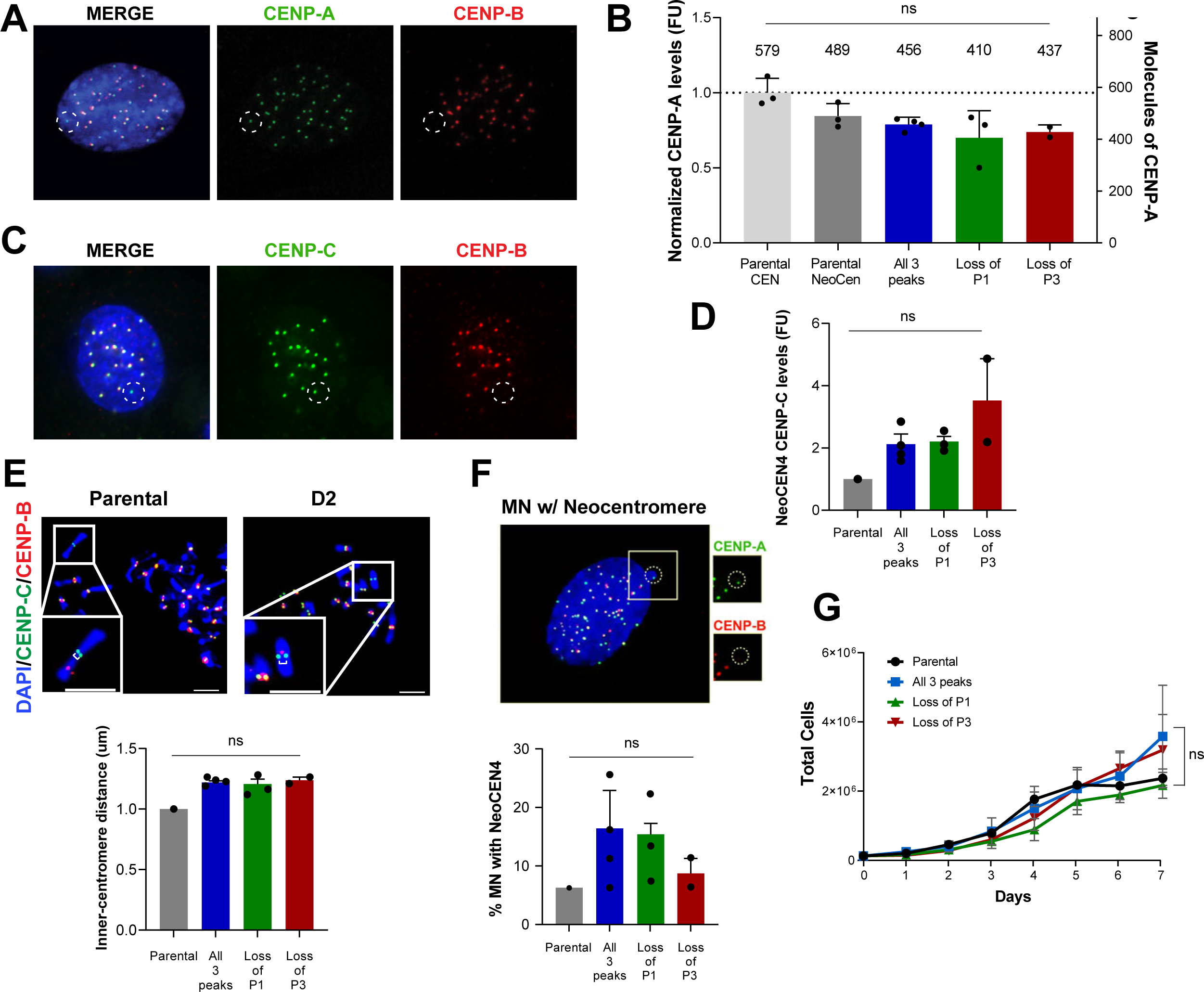
Changes in CENP-A position do not affect centromere function. **A.** NeoCEN4 in parental PD-NC4 cells and clones was identified in interphase IF by locating CENP-A foci lacking CENP-B, which only binds at α-satellite repeats. **B.** Normalized CENP-A fluorescence levels at PD-NC4 parental centromeres, PD-NC4 parental NeoCEN4, and at NeoCEN4 of PD-NC4 clones. Estimated number of CENP-A molecules derived from fluorescent intensity is above each bar. **C.** NeoCEN4 in parental PD-NC4 cells and clones was identified in interphase IF by locating CENP-C foci lacking CENP-B. **D.** Intensity of CENP-C protein at NeoCEN4 in parental PD-NC4 cells and PD-NC4 clones. **E.** Distance between metaphase sister chromatids in parental PD-NC4 cells and clones was calculated by measuring the distance between CENP-C foci at NeoCEN4 (CENP-C not co-localizing with CENP-B) in metaphase spreads. **F.** Micronuclei (MN) in parental PD-NC4 cells and clones characterized as containing NeoCEN4 (+ DAPI, + CENP-A, -CENP-B) were quantified and presented as percentage of MN containing NeoCEN4 out of all MN. **G.** Growth rates of parental PD-NC4 cells and PD-NC4 clones. For panels **B, D, E-G**, PD-NC4 clones were grouped according to CENP-A positional phenotype.

We hypothesized that different CENP-A patterns might affect NeoCEN4 function. Like CENP-A, CENP-C intensity at NeoCEN4 was not significantly different between parental PD-NC4 cells and clones (Fig. 2C, D). We next evaluated sister chromatid cohesion at NeoCEN4 during metaphase but found no significant difference in PD-NC4 clones with different CENP-A positions at NeoCEN4 (Fig. 2E). We quantified micronuclei (MN) containing NeoCEN4 out of total MN but found no significant difference in rates of NeoCEN4 missegregation into MN between clones with different CENP-A positions (Fig. 2F). Lastly, no significant difference was observed in the doubling times of clones with different positions of CENP-A at NeoCEN4 (Fig. 2G). Collectively, these findings suggest that regardless of CENP-A position, a critical number of CENP-A molecules is maintained that preserves neocentromere function.

### CENP-A heterogeneity at human neocentromeres is accompanied with local changes in DNA methylation

We hypothesized that neocentromeres have similar boundary requirements and that different CENP-A positions at NeoCEN4 are accompanied by changes in DNA methylation and/or H3K9me3. To confidently map reads between maternal and paternal NeoCEN4 haplotypes, which exhibit loss of heterozygosity (Fig. S2A-C), we performed Directed Methylation with Long-read sequencing (DiMeLo-seq) (Altemose et al., 2022b) for CENP-A and H3K9me3 in PD-NC4 clones representing the three CENP-A positions at NeoCEN4, aligning the resulting long reads to the recently generated PD-NC4 Chromosome 4 assembly (Fig. 3A, B). We simultaneously used ONT basecalling of native mCpGs to determine DNA methylation status. CENP-A DiMeLo-seq for PD-NC4 clones C2, D2 and E2 confirmed the three distinct CENP-A positions at the active NeoCEN4 locus (paternal/haplotype 1) (Fig. 3D, Fig. S3A). Like conventional centromeres, we find that CENP-A enrichment at NeoCEN4 coincides with dips in DNA methylation and that clonal DNA methylation patterns shifted with CENP-A position. In clone C2, each CENP-A peak correlates with an individual dip in DNA methylation (Fig. 3D). In clones D2 and E2, loss of CENP-A peak 1 or 3 correlates with an increase in DNA methylation at the lost peak position (Fig. 3D). In clone E2, convergence of CENP-A peaks 1 and 2 correlates with a single, wide dip in DNA methylation that encompasses both CENP-A peaks (Fig. 3D). The inactive NeoCEN4 locus on the other copy of chromosome 4 (maternal haplotype/haplotype 2), features 3 regions of hypomethylation roughly correlating to the position of CENP-A at the active NeoCEN4 locus (Fig. 3C, Fig. S4A), suggesting that the existing DNA methylation signature contributed to the distribution pattern of CENP-A established at NeoCEN4.

**Figure 3.**
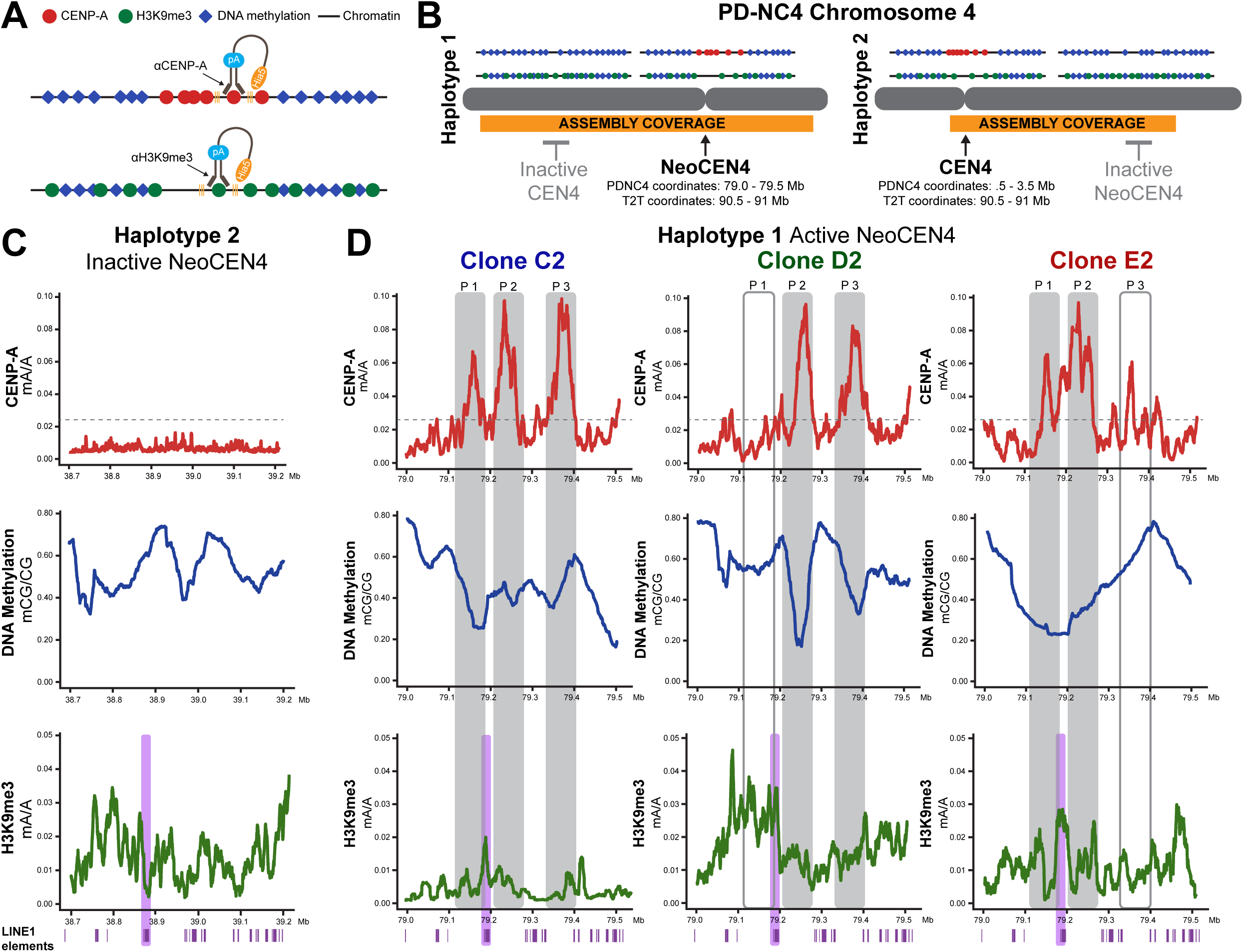
CENP-A heterogeneity at human neocentromeres is accompanied with local changes in DNA methylation. **A.** Experimental schematics of DiMeLo-seq with antibody directed to CENP-A or H3K9me3. **B.** Diagram of PD-NC4 chromosome 4 diploid assembly. Long reads depicted above the assembly highlight ability to assess DNA methylation and H3K9me3 at loci that have lost or gained CENP-A and compare to the matched locus in the other chromosome 4 haplotype. The NeoCEN4 and CEN4 coordinates are different within the PD-NC4 assembly and the T2T CHM13v2 assembly. **C.** DiMeLo-seq at the inactive NeoCEN4 locus on chromosome 4 haplotype 2 in PD-NC4 clone D2. Purple bars highlight position of LINE1 element refractive to CENP-A acquisition. **D.** DiMeLo-seq at the active NeoCEN4 locus on chromosome 4 haplotype 1 in PD-NC4 clones representing the three CENP-A positional phenotypes: three CENP-A peaks (clone C2), loss of CENP-A peak 1 (clone D2) and loss of CENP-A peak 3 (clone E2). Horizontal grey dashed line indicates threshold for CENP-A enrichment. Grey shadows indicate the position of CENP-A peaks in parental PD-NC4 cells. Purple bars highlight position of LINE1 element refractive to CENP-A acquisition. In **C-D**, DiMeLo-seq tracks in red (CENP-A), blue (DNA methylation), and green (H3K9me3) depict the average fraction of modified bases over 5000 bp windows with probability ≥ 0.8.

We found H3K9me3 to be present at the inactive locus of NeoCEN4 at variably low-moderate levels (Fig. 3C, Fig. S4A). At the active NeoCEN4 locus H3K9me3 levels were also variable between clones, but increased in clone D2 at the peak 1 position where CENP-A is lost, confirming previous observations of anti-correlation between CENP-A and H3K9me3 (Martins et al., 2016; Ohzeki et al., 2012). Notably, the active NeoCEN4 locus gained H3K9me3 in all clones at a 6 kb L1HS LINE element found between CENP-A peak 1 and peak 2 that is not enriched for H3K9me3 in the inactive NeoCEN4 locus (Fig. 3C, D, Fig. S4A). LINEs and other transposable elements are often enriched in pericentromeric heterochromatin where their silencing may play a role in boundary maintenance (Hartley and O’Neill, 2019). We observed variable H3K9me3 in the 7 Mb of chromatin surrounding NeoCEN4, and could not detect a shared pattern for H3K9me3 at either the active or inactive NeoCEN4 locus among clones, highlighting their epigenetic heterogeneity (Fig. 3C, D, Fig. S3C-D, S4A).

### Human native repetitive centromeres are epigenetically heterogenous but are confined within a single wide CDR

The active PD-NC4 CEN4 (maternal/haplotype 2) recruits CENP-A to HOR1 and DiMeLo-seq reveals a prominent CENP-A peak at coordinates ∼1.45 to 1.65 Mb in HOR1 that correlates with a wide dip in DNA methylation and a dip in H3K9me3 (Fig. 4A, B, Fig. S3B, S5A). A close examination of CENP-A at the active CEN4 reveals CENP-A enrichment ranging from ∼150 - 170 kb in clones, with a shared highly enriched 60 kb CENP-A peak at coordinates 1.48 to 1.54 Mb (Fig. 4C). Outside of that peak, CENP-A position and enrichment varied among clones (Fig. 4C). The heterogeneously enriched CENP-A region at active CEN4 was confined within a single wide CDR ranging in size between 260 - 320 kb in clones (Fig. 4C). H3K9me3 pattern varied among PD-NC4 clones at the active CEN4, with only clone D2 showing a clear dip in H3K9me3 that correlated with the position of the conserved CENP-A peak (Fig. 4C), mirroring the heterogeneity in H3K9me3 we observed at NeoCEN4. DiMeLo-seq revealed no CENP-A enrichment nor CDR in the heavily methylated inactive CEN4 (paternal/haplotype 1) (Fig. 4C, Fig. S5A). H3K9me3 at the inactive CEN4 was uniformly moderate across HOR1 and did not decrease as observed at the site of the conserved CENP-A peak in the active CEN4 haplotype (Fig. 4A, Fig. S5A).

**Figure 4.**
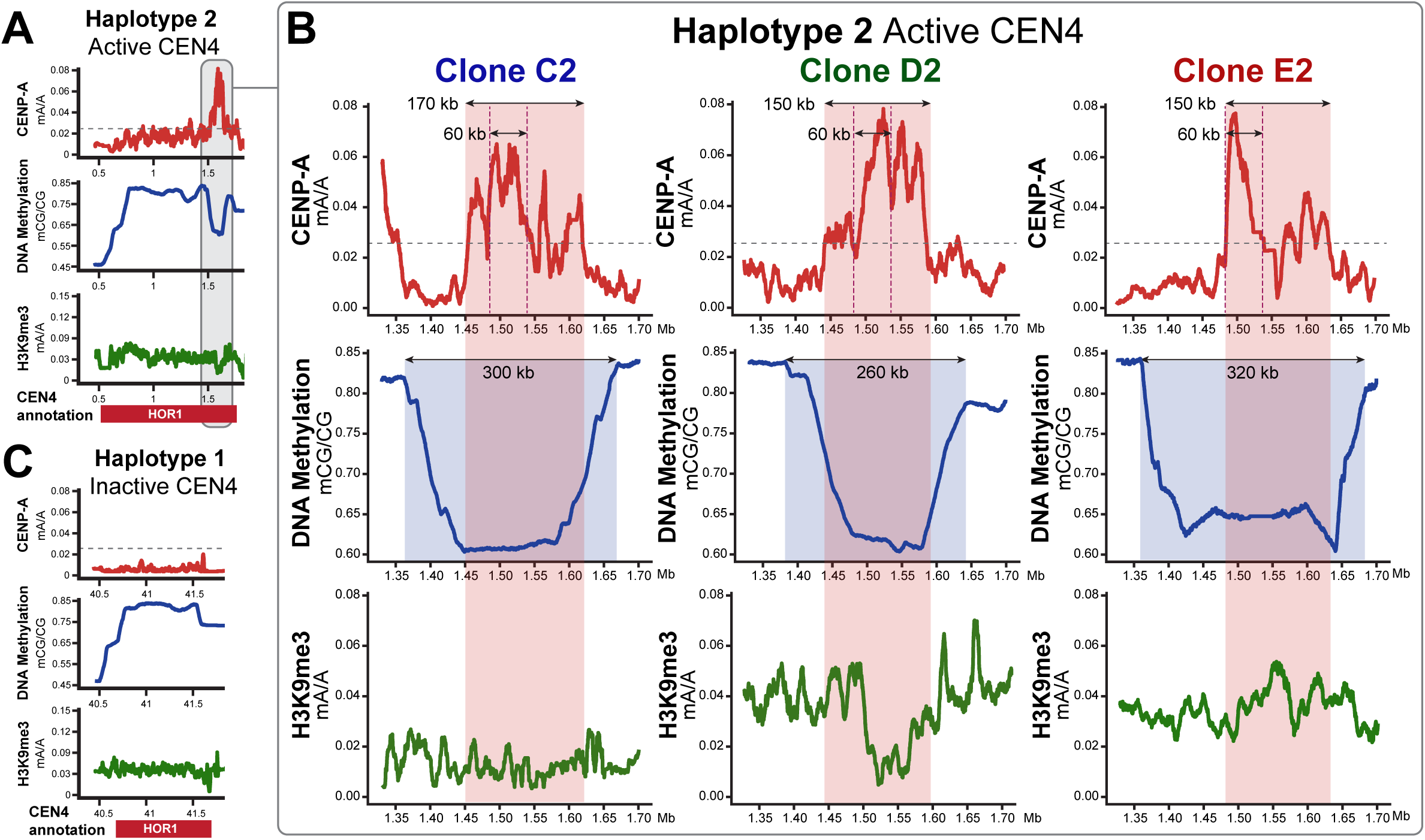
Human native repetitive centromeres are epigenetically heterogenous but are confined within a single wide CDR. **A.** DiMeLo-seq at the active CEN4 locus in clone D2. **B.** Expanded view of DiMeLo-seq at the region of active CEN4 locus where kinetochore is likely to assemble (grey shadow in **A**) from each PD-NC4 clone shown in Fig. 3D. Horizontal grey dashed line indicates threshold for CENP-A enrichment. Vertical red dashed line indicates the position of a conserved CENP-A peak found in each clone. Red highlight indicates the size of CENP-A enriched area in each clone. Blue highlight indicates the size of CDR in each clone. **C.** DiMeLo-seq at the inactive CEN4 locus in clone D2. DiMeLo-seq tracks in red (CENP-A), blue (DNA methylation), and green (H3K9me3) depict the average fraction of modified bases over 5000 bp windows with probability ≥ 0.8.

### Prolonged cellular proliferation accelerates NeoCEN4 drift, evolution and instability through loss of DNA methylation CDRs

To evaluate long-term stability of CENP-A position, we maintained single cell-derived PD-NC4 clones in culture over 12 months (PD-NC4 doubling time 36h; ∼280 cell cycles). Only clone C4 did not maintain NeoCEN4 over 12 months (Fig. S1A, Fig. S5B, Table S1). PD-NC4 clones that lost CENP-A peaks 1 or 3 did not re-acquire it over 12 months (Fig. 5A-C, Fig. S5B) and two clones (clone A1 and C3) lost a CENP-A peak within 6 months (Fig. S5B). Importantly, CENP-A position at NeoCEN4 naturally evolved over prolonged proliferation and exhibited large-scale (5 – 42 kb) positional drift/expansion/shrinkage relative to parental PD-NC4 cells (Fig. 5A-C, Fig. S5B). CENP-A remained conspicuously absent in all clones from the 6 kb L1HS LINE element between peak 1 and peak 2 (Fig. 5A-C, Fig. S5B) that gained H3K9me3 in the active NeoCEN4 haplotype (Fig. 3C, D, Fig. S4A-B). Indeed, we find LINE1 elements distributed across the NeoCEN4 locus appear to restrict CENP-A position, suggesting LINE1 elements serve as neocentromere boundaries (Fig. 5A-C, Fig. S5B), as previously proposed for native centromeres (Hoyt et al., 2022). Accordingly, we observe a propensity of CENP-A peak 1 to dramatically drift/expand upstream, consistent with decreased LINE1 element distribution at this region of NeoCEN4 (Fig. 5A-C). Notably, we never observed loss of CENP-A peak 2 or concurrent loss of CENP-A peaks 1 and 3 in any clone at any timepoint, indicating that maintaining central peak 2 and either peak 1 or 3 is critical for NeoCEN4 function.

**Figure 5.**
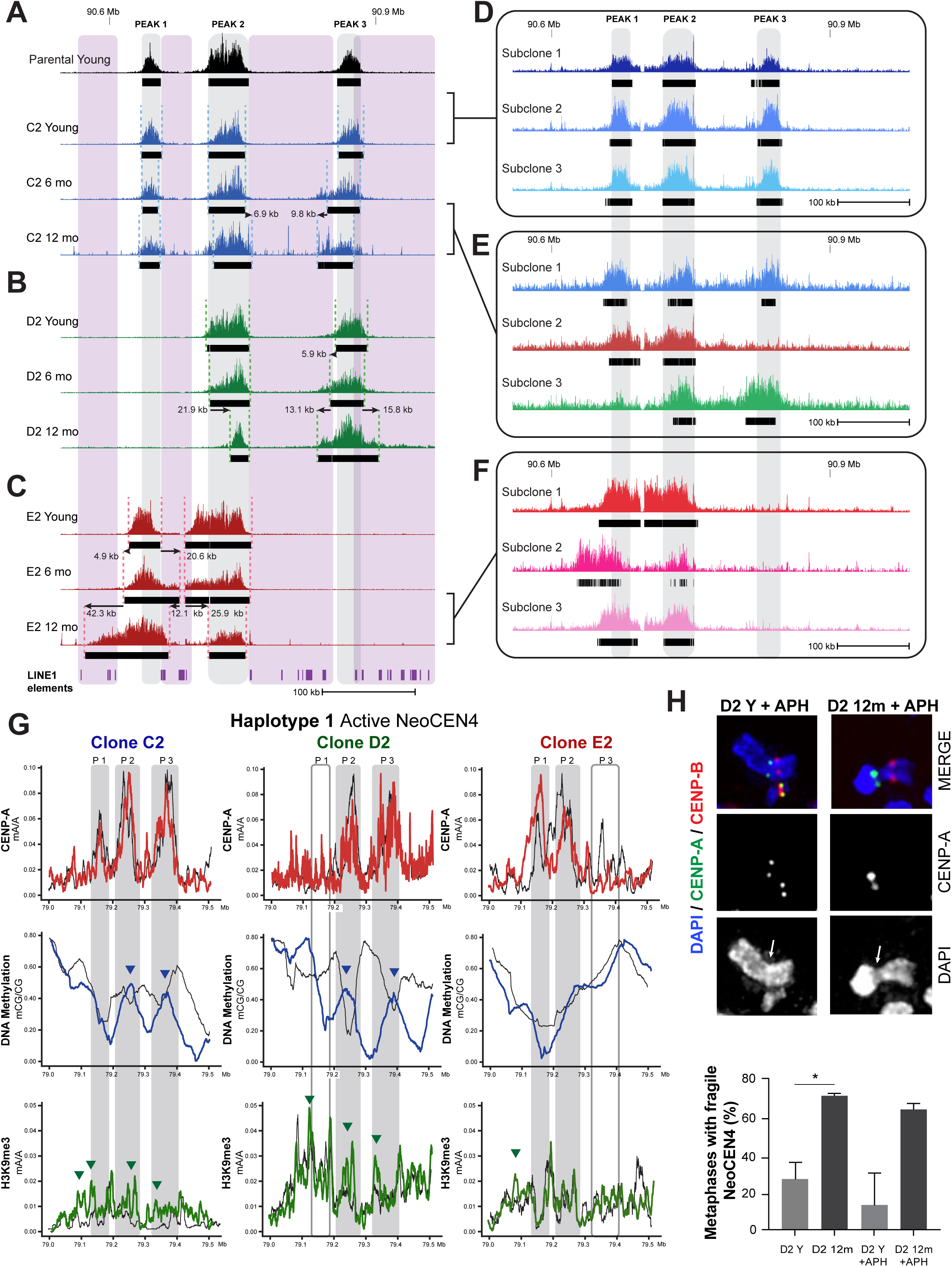
Prolonged cellular proliferation accelerates NeoCEN4 drift, evolution and instability through loss of DNA methylation CDRs. **A.** CENP-A ChIP-seq or CUT&RUN for young PD-NC4 parental cells and clone C2 (three CENP-A peaks) when young and after 6 and 12 months in culture. **B.** CENP-A ChIP-seq or CUT&RUN for clone D2 (loss of CENP-A peak 1) when young and after 6 and 12 months in culture. **C.** CENP-A ChIP-seq or CUT&RUN for clone E2 (loss of CENP-A peak 3) when young and after 6 and 12 months in culture. **D.** CENP-A CUT&RUN for subclones isolated from clone C2 when young. **E.** CENP-A CUT&RUN for subclones isolated from clone C2 after 12 months in culture. **F.** CENP-A CUT&RUN for subclones isolated from clone E2 after 12 months in culture. For **A-F**, grey shadows represent the position of CENP-A peaks in young parental PD-NC4 cells. Black bars show location of significantly enriched CENP-A peaks (p<0.00001, fc ≥ 20). For **A, B** and **C**, dotted lines mark the position of significantly enriched CENP-A peaks in clones at each timepoint. Arrows indicate movement of significantly enriched CENP-A peaks in clones relative to their earlier timepoint. Purple shadows mark areas enriched for LINE1 elements in PD-NC4 cells. **G.** DiMeLo-seq after 12 months at the active NeoCEN4 locus on chromosome 4 haplotype 1 in PD-NC4 clones C2 (three peaks), D2 (loss of peak 1) and E2 (loss of peak 3). Grey shadows indicate the position of CENP-A peaks in parental young PD-NC4 cells. Blue arrows highlight where DNA methylation dips have become peaks after 12 months. Green arrows highlight positions where H3K9me3 levels have increased after 12 months. DiMeLo-seq tracks in red (CENP-A), blue (DNA methylation), and green (H3K9me3) depict the average fraction of modified bases over 5000 bp windows with probability ≥ 0.8, and are overlayed on matched DiMeLo-seq tracks at young timepoint that are depicted in black. **H.** Immunofluorescence of CENP-A (green) and CENP-B (red) to detect fragility of NeoCEN4 region in PD-NC4 clone D2 with and without treatment with aphidicolin at young and 12 months timepoints. Quantification (below) of NeoCEN4 fragility in D2 Y and D2 12m cells without and with aphidicolin (+APH). Error bars are standard deviation; p<0.05, two-tailed t-test.

We hypothesized that the significant expansion in CENP-A peak 1 of clone E2 after 12 months (Fig. 5C) represents an average within the E2 12 months population. Subcloning of the E2 12 months population followed by CENP-A CUT&RUN revealed acquired heterogeneity in CENP-A position at NeoCEN4, with two subclones maintaining CENP-A position similar to E2 6 months population (Fig. 5F, subclones 1 and 3) and one subclone experiencing significant drift and expansion of peak 1, as well as shrinkage of peak 2 (Fig. 5F, subclone 2). Thus, CENP-A position at NeoCEN4 of E2 12 months represents a population average. To determine when heterogeneity in CENP-A position is acquired, we isolated subclones from clone C2 at young (∼40 cell cycles following single-cell sub-cloning) and 12 months timepoints (∼280 cell cycles). While subclones of C2 young population had nearly identical CENP-A positions (Fig. 5D), subclones of C2 12 months population acquired the heterogeneity of parental PD-NC4 cells, exhibiting three distinct CENP-A positions (Fig. 5E). Taken together, our findings indicate that CENP-A position is stably maintained at NeoCEN4 for at least ∼40 cell cycles, but not over prolonged cellular proliferation.

Next, we performed DiMeLo-seq for CENP-A and H3K9me3 in the same PD-NC4 clones after 12 months to evaluate changes in DNA methylation and H3K9me3 at NeoCEN4 over prolonged cellular proliferation. At the inactive NeoCEN4 locus, CENP-A remained absent, overall DNA methylation levels were increased, and H3K9me3 retained largely the same pattern (Fig. S4B-C). At the active NeoCEN4 locus, CENP-A DiMeLo-seq confirmed changes observed with CUT&RUN in CENP-A position in 12 months old clones (Fig. 5G). Unexpectedly, we observed local increase in DNA methylation at NeoCEN4 regions enriched for CENP-A in C2 and D2 cells after 12 months that indicated decoupling of CENP-A peaks from their individual CDRs (Fig. 5G). Since this observation might be obscured by the significant heterogeneity acquired at NeoCEN4 after 12 months (Fig. 5E, F), we performed DiMeLo-seq on subclone 1 from C2 12 months population (Fig. 5E) and confirmed that after prolonged cellular proliferation CENP-A enrichment is no longer correlated with CDR placement at NeoCEN4 (Fig. S4D). H3K9me3 levels increased at regions enriched for CENP-A that lost correlation with CDRs in clones C2 and D2 at 12 months (Fig. 5G, arrows). In contrast, in clone E2 that had a wider CDR when young, CENP-A position remained largely associated with the CDR after 12 months and H3K9me3 levels were not increased within CENP-A peaks (Fig. 5G). To test whether prolonged proliferation-induced loss of DNA methylation CDRs and gain of H3K9me3 at regions enriched for CENP-A affect NeoCEN4 function, we treated clone D2 young and 12 months old cells with aphidicolin to induce replication stress, then quantified NeoCEN4 fragility (Fig. 5H). Prolonged proliferation increased fragility of NeoCEN4 2.65-fold without replication stress in untreated cells, and 5.15-fold in aphidicolin treated D2 12 months cells relative to aphidicolin treated D2 young cells (Fig. 5H). Taken together, our results demonstrate that loss of DNA methylation CDRs and gain of H3K9me3 at CENP-A peaks after 12 months drive NeoCEN4 instability and fragility.

### Native centromeres maintain a single wide CDR over prolonged cellular proliferation

Next, we used DiMeLo-seq to determine CENP-A, DNA methylation and H3K9me3 occupancy at PD-NC4 CEN4 over 12 months of cellular proliferation (Fig. 6A). Similarly to NeoCEN4, we found CEN4 to evolve epigenetically over 12 months. While we found that the position of the conserved CENP-A peak present in young clones was precisely maintained, we observed changes in CENP-A position including expansion of 65 kb and 145 kb in clones D2 and E2, respectively (Fig. 6B), indicating acquisition of CENP-A positional heterogeneity at CEN4, as seen at NeoCEN4 (Fig. 5E-F). Changes were also evident between the CDRs of young and 12 months old clones, including contraction of 20 and 27 kb in clone C2 and E2 respectively, and expansion of 75 kb in clone D2 (Fig. 6B). Despite these changes in CENP-A position and CDR size over 12 months, the single CDR remained at the same overall position in HOR1 and CENP-A remained confined within the boundaries of the CDR (Fig. 6A, B). We next quantified the variation in CENP-A position between individual DiMeLo-seq reads mapped to CEN4 in C2 young and 12 months populations and found that position of CENP-A acquires increased heterogeneity within the C2 population over 12 months, while CENP-A position in C2 young is maintained with precision and low variation (Fig. S5C-D). Importantly, H3K9me3 patterns and levels remained relatively unchanged/similar at CEN4 over the course of 12 months (Fig. 6B). These findings strongly demonstrate that native, repetitive human centromeres also naturally evolve epigenetically over prolonged cellular proliferation. A key distinction between CEN4 and NeoCEN4 is the maintenance of CENP-A within the stable boundaries of a single wide CDR and H3K9me3 stability at CEN4 over 12 months, demonstrating that native centromeres have greater epigenetic stability than neocentromeres over prolonged cellular proliferation.

**Figure 6.**
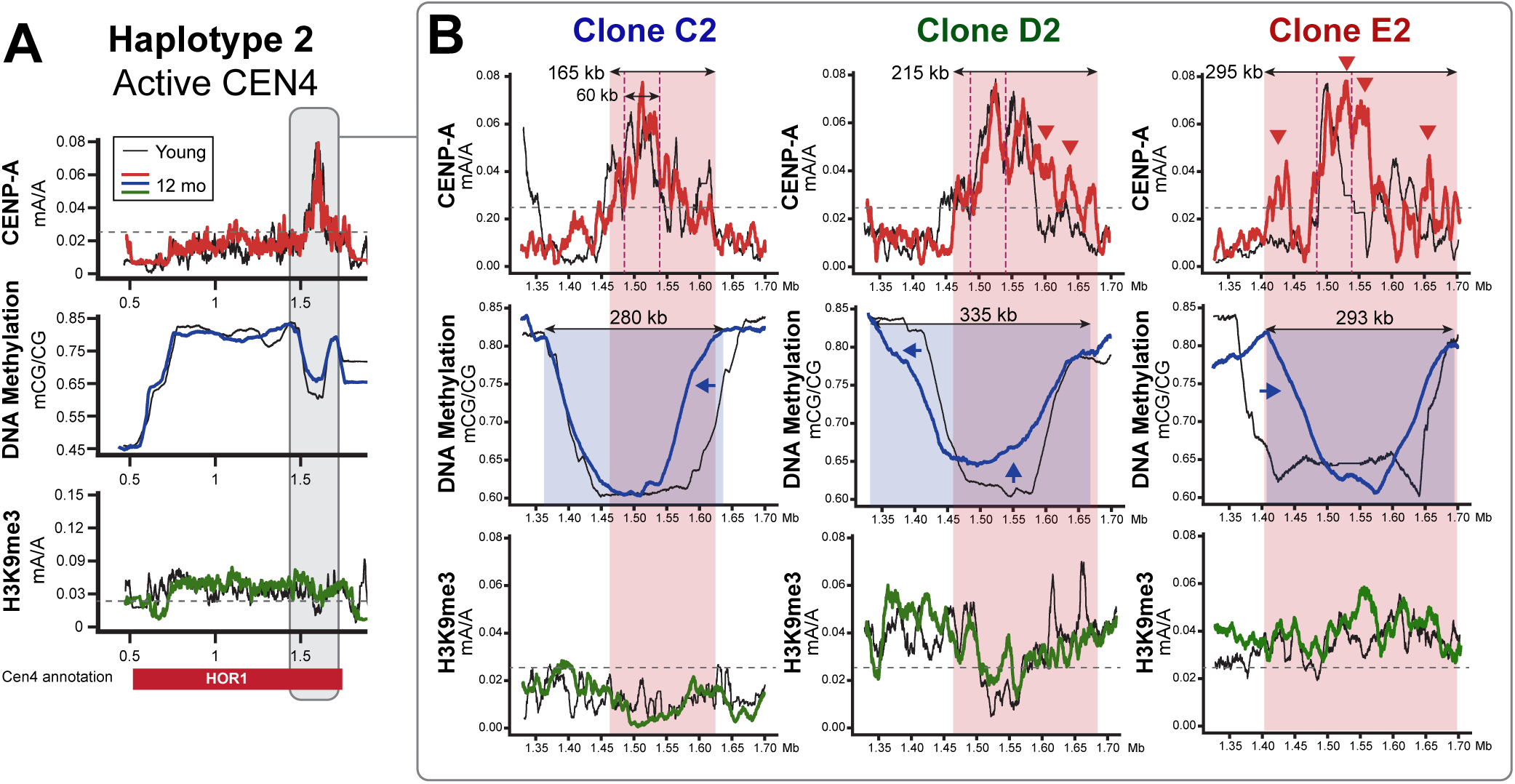
Native centromeres maintain a single wide CDR over prolonged cellular proliferation. **A.** DiMeLo-seq at the active CEN4 locus in clone D2 after 12 months. **B.** Expanded view of DiMeLo-seq after 12 months at the region of active CEN4 locus where kinetochore is likely to assemble (grey shadow in **A**) from PD-NC4 clones C2 (three peaks), D2 (loss of peak 1) and E2 (loss of peak 3). Horizontal grey dashed line indicates threshold for CENP-A enrichment. Vertical red dashed line indicates the position of a conserved CENP-A peak found in each clone. Red highlight indicates the size of CENP-A enriched area in each clone. Blue highlight indicates the size of CDR in each clone. Blue arrows highlight changes in the CDR after 12 months. DiMeLo-seq tracks in red (CENP-A), blue (DNA methylation), and green (H3K9me3) depict the average fraction of modified bases over 5000 bp windows with probability ≥ 0.8, and are overlayed on DiMeLo-seq tracks at young timepoint that are depicted in black.

### Prolonged CENP-A and HJURP overexpression destabilizes NeoCEN4 and drives neocentromere re-localization

The striking heterogeneity and plasticity of CENP-A and DNA methylation at NeoCEN4 led us to predict that overexpression of CENP-A and HJURP would lead to increased deposition and incorporation of CENP-A at this site, causing increased drift and/or expansion of NeoCEN4. To test this hypothesis, we transduced PD-NC4 cells with lentiviral vectors expressing CENP-A^LAP-eYFP^ and HJURP^LAP-mCherry^, selected single-cell derived clones with various levels of CENP-A and HJURP overexpression and maintained those in culture for 12 months (Fig. 7A, Fig. S6A-C). Using CENP-A CUT&RUN we first examined native CEN4 and found that in clone G1 that overexpresses CENP-A and HJURP at low levels (7x/4x, respectively), CENP-A enrichment increased 69% compared to non-overexpressing PD-NC4 cells across HOR1 and 35% within the active region of CEN4 and remained similarly enriched at 6 and 12 months (Fig. 7B-C, Fig. S6D-E), indicating increased CENP-A binding at HOR1 of CEN4. Increasing CENP-A/HJURP overexpression to 9x/4x (clone E3) and 47x/13x (clone C4) resulted in reduction of total area enriched for CENP-A and number of CENP-A reads mapped at CEN4 that continued to decline with time (Fig. 7B-C, Fig. S6D-E), indicating that increased CENP-A and HJURP overexpression leads to gradual depletion of CENP-A at CEN4 which may be linked to increased ectopic CENP-A deposition (Lacoste et al., 2014; Mahlke and Nechemia-Arbely, 2020; Nechemia-Arbely et al., 2019; Shrestha et al., 2017; Shrestha et al., 2021).

**Figure 7.**
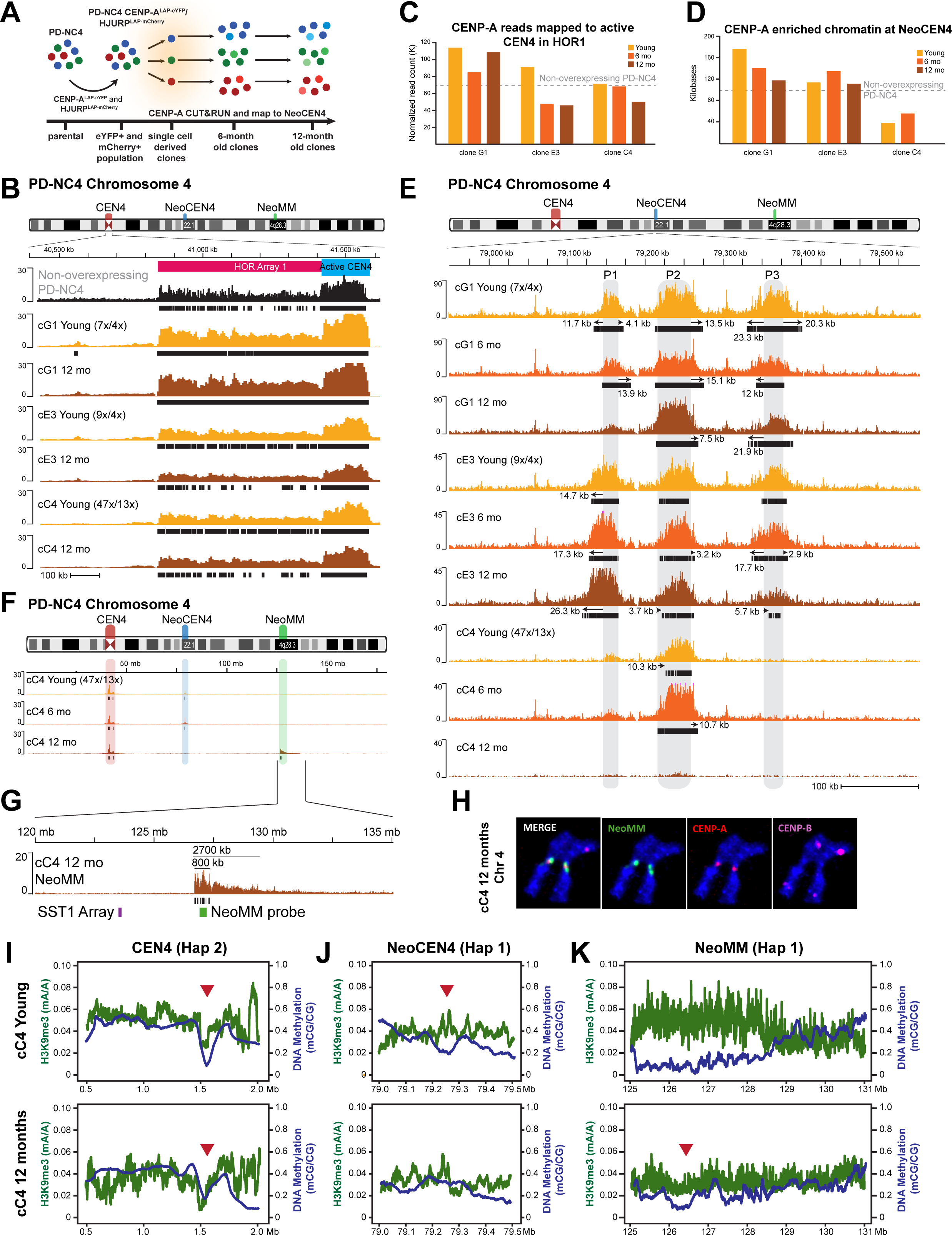
Prolonged CENP-A and HJURP overexpression destabilizes NeoCEN4 and drives neocentromere re-localization. **A**. Experimental design for deriving CENP-A/HJURP overexpressing PD-NC4 clones and assessing effect on CENP-A position across 12 months. **B.** CENP-A CUT&RUN for non-overexpressing and CENP-A/HJURP overexpressing PD-NC4 clones aligned to PD-NC4 chromosome 4 assembly at young and 12 months timepoints at CEN4. Colored bar above tracks indicates positions of HOR1 and active centromere region at CEN4 HOR1. **C.** Quantification of CENP-A-bound reads aligning to the active centromere region at CEN4 in CENP-A/HJURP overexpressing PD-NC4 clones over 12 months. **D.** Quantification of CENP-A enriched chromatin at NeoCEN4 in CENP-A/HJURP overexpressing PD-NC4 clones over 12 months. In **C** and **D**, grey dashed line indicates average level in non-overexpressing PD-NC4 clones (n=3). **E.** CENP-A CUT&RUN at NeoCEN4 in CENP-A/HJURP overexpressing PD-NC4 clones at young, 6 months, and 12 months timepoints. Grey shadows indicate positions of significantly enriched CENP-A peaks in young parental PD-NC4 cells. **F.** CENP-A CUT&RUN in CENP-A/HJURP overexpressing clone C4 over 12 months. **G.** CENP-A CUT&RUN at NeoMM in clone C4 after 12 months in culture. Purple bar indicates position of the repetitive SST1 array in band 4q28.3 and location of a known fragile site. Green bar indicates position of NeoMM FISH probe. In **B, E,** and **F** Chromosome ideogram at top features red, blue and green shadows highlighting position of CEN4, NeoCEN4, and NeoMM, respectively. In **B, E, F,** and **G,** black bars show location of significantly enriched CENP-A peaks (p<0.00001, fc ≥ 20). **H.** IF-FISH on metaphase spread in clone C4 12months cells with CENP-A (red), CENP-B (magenta) and NeoMM probe (green) confirms CENP-A localization at NeoMM locus. Genomic location of NeoMM probe is depicted in **(G). I, J, K.** DiMeLo-seq at active CEN4 **(I)**, NeoCEN4 **(J)**, and NeoMM **(K)** in CENP-A/HJURP overexpressing clone C4 when young (top) and after 12 months (bottom). DiMeLo-seq tracks in green (H3K9me3) and blue (DNA methylation) depict the average fraction of modified bases over 5000 bp windows with probability ≥ 0.8. Red arrows represent the position of CENP-A enrichment from CUT&RUN in **B, E,** and **G.**

At NeoCEN4, while CENP-A and HJURP overexpression did not accelerate CENP-A drift from the positions observed in parental PD-NC4 cells, we observed a 109% expansion in total CENP-A enrichment, up to 175 kb, compared to 84 kb in non-overexpressing parental PD-NC4 cells (Fig. 7D, E, compare to Figs. 1, 4; Table S4). Over prolonged cellular proliferation, clone G1 showed gradual loss of Peak 1 (Fig. 7E), similarly to non-overexpressing clone A1 (Fig. S5B), and clone E3 exhibited gradual shrinkage of CENP-A peak 3 (Fig. 7E), similar to non-overexpressing clone C3 (Fig. S5B). Conversely, Clone C4 that had high CENP-A and HJURP overexpression (47x and 13x, respectively) lost both CENP-A peaks 1 and 3 (Fig. 7E), a pattern never observed in non-overexpressing clones (Figs. 1, 5; Table S4). In clone C4, NeoCEN4 spans only 34.2 kb in young and attempts to compensate by expanding to 50.3 kb after 6 months (Fig. 7D, E). After 12 months of high CENP-A and HJURP overexpression, CENP-A was completely lost from NeoCEN4 in clone C4 (Fig. 7E-F) and a new neocentromere, termed NeoMM, emerged ∼50 Mb away in the q arm of chromosome 4 (Fig. 7F-H). NeoMM formed at 4q28.3 ∼3 Mb downstream from a known fragile site (Tremblay et al., 2010) that contains an SST1 array composed of large, 2.4 kb monomers (Fig. 7G). NeoMM spans over a large region of ∼2.7 Mb with the highest enrichment for CENP-A occurring in an ∼800 kb region. IF-FISH with a 190 kb probe within the highly enriched CENP-A region confirmed that CENP-A has relocated to the NeoMM locus and indicated that Chr4 has formed an isochromosome with two ancestral inactive centromeres marked by CENP-B flanking NeoMM, that now functions as the new active centromere (Fig. 7H). To investigate how centromere epigenetic heterochromatin signatures may have changed in response to prolonged CENP-A/HJURP overexpression and/or NeoCEN4 re-localization, we performed DiMeLo-seq for H3K9me3 and DNA methylation in clone C4 when young and after 12 months of CENP-A/HJURP overexpression. At native CEN4, prolonged overexpression of CENP-A/HJURP had little effect on DNA methylation or H3K9me3 (Fig. 7I; compare to Fig. 6A). At NeoCEN4, presence of the CDRs decreased in accordance with CENP-A loss upon elevated CENP-A/HJURP overexpression. The single remaining significantly enriched CENP-A peak in clone cC4 young (Fig. 7E) was accompanied by a single corresponding CDR (Fig. 7J; compare to Fig. 3D). The loss of CDR at NeoCEN4 became more pronounced after 12 months when CENP-A completely vacated NeoCEN4 (Fig. 7E, J). Interestingly, we find that the emerging neocentromere, NeoMM, formed at a locus with relatively high H3K9me3 but low DNA methylation levels (Fig. 7K, top), suggesting again that low DNA methylation serves as a permissive environment for CENP-A seeding (Fig. S4A). After 12 months of elevated CENP-A/HJURP overexpression and CENP-A re-localization to the NeoMM locus, we find that this region has been epigenetically remodeled to reduce H3K9me3 and form DNA methylation CDR-like regions that encapsulate CENP-A (Fig. 7K, bottom). Our findings implicate CENP-A and HJURP overexpression in centromere and neocentromere destabilization over prolonged cellular proliferation and demonstrate that overexpression of CENP-A and HJURP can drive neocentromere formation and reshape local epigenetics.

## Discussion

Precise positional maintenance of CENP-A helps prevent centromeres from forming in more than one location on a chromosome (Mahlke and Nechemia-Arbely, 2020; Nechemia-Arbely et al., 2019), which could lead to errors in chromosome segregation during mitosis. Indeed, CENP-A has been observed as a stable, long-lived nucleosome that is inherited indefinitely and precisely to maintain and propagate centromere identity (Fachinetti et al., 2013; Mahlke et al., 2023; Nechemia-Arbely et al., 2019; Ross et al., 2016; Smoak et al., 2016). Nevertheless, our study of three human neocentromeres and a native, repetitive human centromere is the first to report that CENP-A position and patterns of DNA methylation and H3K9me3 are heterogenous at the same centromere of single cells within a cell population (Figs. 1, 3D, and 4C), and these epigenetically distinct centromeres are equally functional (Fig. 2). Thus, epigenetic plasticity is a general feature of a single human centromere or neocentromere within an identical cell population (Fig. S7).

Our work uses DiMeLo-seq to assess epigenetic modifications on long reads and align long reads to a cell line-matched assembly, ensuring high fidelity recapitulation of the centromere and neocentromere epigenetic landscape on both haplotypes of the PD-NC4 chromosome 4. Importantly, our finding of centromere epigenetic plasticity between single cells of human cell populations has implications for how we interpret centromere epigenetics sampled from a group of cells, such as ChIP-seq or CUT&RUN, and highlights that these analyses may mask divergent single-cell level patterns. This newly found single-cell level centromere epigenetic plasticity is key to understanding how rapidly evolving human centromeres that have acquired distinct, lineage-dependent HOR array organization at the same centromere are capable of maintaining one functional centromere on each chromosome (Altemose et al., 2022a; Logsdon et al., 2024; Miga and Alexandrov, 2021).

Our study is also the first to report the presence of DNA methylation CDRs at neocentromeres. The presence of three CDRs at the inactive NeoCEN4 locus suggests it was initially formed by CENP-A deposition into a permissive epigenetic environment, where it was constrained by the existing DNA methylation pattern as well as the presence of flanking LINE1 elements, which are often repressed by DNA methylation and H3K9 methylation (Klein and O’Neill, 2018). We find that CENP-A is precisely maintained within CDRs at CEN4 and NeoCEN4 (Figs. 5D, S7) for at least ∼40 cell cycles, confirming our previous conclusions of precise CENP-A maintenance during DNA replication (Nechemia-Arbely et al., 2019). On average, human cells divide 40-60 times during development (Hayflick, 1965), suggesting that for most human cells CENP-A position will be precisely maintained throughout their lifespan. Over prolonged cellular proliferation (∼280 cell cycles, 12 months) CENP-A at CEN4 and NeoCEN4 acquires significant positional heterogeneity (Fig. 5E, F; Fig. S7). At NeoCEN4, CENP-A is no longer constrained by LINE1 elements, fails to remain strictly associated with a CDR and some of its peaks gain H3K9me3 (Fig. 5G; Fig. S4D, Fig. S7), which has been shown to be incompatible with CENP-A chromatin and to contribute to neocentromere instability (Murillo-Pineda et al., 2021; Ohzeki et al., 2016; Ohzeki et al., 2012). At CEN4, variable CENP-A position occurs within the boundaries of a single wide CDR that remains stable over 12 months with stable H3K9me3 pattern, indicating that native human centromeres have greater long-term epigenetic stability than human neocentromeres (Fig. S7). Collectively, these results suggest that inherent epigenetic instability underlies functional shortcomings of neocentromeres (Bassett et al., 2010; Fachinetti et al., 2015) and highlight the roles of the CDR and H3K9me3 in maintaining the long-term epigenetic stability of native centromeres, possibly through CENP-B and DNA methylation interaction at α-satellite sequences (Tanaka et al., 2005).

Regardless of changes in CENP-A position, NeoCEN4 maintains a threshold above the 400 CENP-A molecules expected at a normal repetitive human centromere (Bodor et al., 2014), indicating that specific CENP-A position is less important than preserving a local concentration of CENP-A sufficient for centromere function. Overexpression of CENP-A and HJURP at high levels caused CENP-A depletion from NeoCEN4, prompting the formation of a new neocentromere (NeoMM) in the q arm of chromosome 4 (Fig. 7). Thus, balanced CENP-A and HJURP expression levels play a vital role in preserving the position and stability of epigenetically plastic and evolving centromeres and neocentromeres (Fig. S7). The emergence of NeoMM in a region with high H3K9me3 and low DNA methylation, the establishment of CDR-like methylation dips at NeoMM upon CENP-A relocation, the shifting CDRs in accordance with CENP-A position at NeoCEN4, and the loss of CDRs at NeoCEN4 upon re-localization of CENP-A to NeoMM collectively suggest that DNA methylation plays a more important boundary role at centromeres and neocentromeres than does H3K9me3.

Strict positional maintenance of CENP-A within rapidly evolving centromeric DNA provides opportunities for centromeres to be fractured or lost through evolutionary accumulation of mutations and rearrangements. Instead, we find the heterogenous and plastic epigenetic landscape of human centromeres within a population of cells allows for flexible CENP-A maintenance within the naturally evolving boundaries of the CDR and flanking H3K9me3, preserving a single functional CENP-A domain within rapidly evolving centromeric DNA (Fig. S7). Given the number of cell cycles (∼280) we observed necessary for dramatic changes to centromere epigenetics, our findings suggest that cell types with high mitotic indexes such as gut, skin epithelium and tumor cells are most likely to be impacted by centromere epigenetic plasticity. Further, the occurrence of aneuploidy and frequent emergence of neocentromeres in human cancers indicate a possible connection between centromere epigenetic plasticity and tumor heterogeneity.

## Acknowledgments

We thank Dr. Ofer Shoshani (Weizmann Institute) and Dr. Peter Ly (UT Southwestern) for valuable discussions, Dr. Peter Warburton for essential cell lines. We thank members of our laboratories for their valuable suggestions during the manuscript writing. We thank Danilo Dubocanin and Glennis Logsdon for technical advice during bioinformatic analysis. This work was supported by the US National Institutes of Health (Grant no. R35 GM142717 to Y.N.A, Grant no. R01 HG009190 to W.T., and Grant no. R01GM123312 to R.J.O.), and by a Searle Scholars Program award to K.H.M. Work in the Altemose Lab was supported by a Howard Hughes Medical Institute Hanna H. Gray Faculty Fellowship and by a Chan Zuckerberg Biohub Investigatorship.

This research was supported in part by the National Cancer Institute of the National Institutes of Health (NIH Award Number P30CA047904), the UPMC Hillman Cancer Center Cytometry Facility (RRID:SCR_025361) and by the University of Pittsburgh Center for Research Computing HTC cluster (NIH Award Number S10OD028483). The funders had no role in study design, data collection, analysis, publication decision, or manuscript preparation.

## Author Information

These authors contributed equally: Cy Chittenden, Savannah Hoyt, and Jonas Koeppel.

## Contributions

M.A.M., Y.N.A. conceived the project, designed the experiments, and wrote the manuscript. M.A.M., L.L., P.N., R.R., K.Z., conducted the experiments and analyzed the data. J.K. and P.J.C. analyzed whole genome sequencing experiments. Y.X., C.C., K.H.M. and N.A. performed DiMeLo-seq experiments and provided training on DiMeLo-seq methods and analysis. S.H. and R.J.O. assembled reference assembly for PD-NC4 chromosome 4. P.W.H. and W.T. provided training on ONT long reads sequencing. All authors provided input on the manuscript.

## Supplemental Figure Legends

**Figure S1. NeoCEN4 and NeoCEN13 are maintained within their respective populations, related to Figure 1. A.** Representative metaphase spreads of PD-NC4 parental cells and single-cell derived clones at young, 6 months, and 12 months timepoints. **B.** Representative metaphase spreads of IMS13q parental population and single-cell derived clones. **C**. Representative metaphase spreads of BBB parental population and clones and BBB parental cells and single-cell derived clones. In **A-C**, insets highlight the neocentromere where CENP-A (green) does not co-localize with CENP-B (red) that marks α-satellite containing native centromeres. **D.** Whole genome sequencing of parental PD-NC4 cells and clones C2 (3 CENP-A peaks), D2 (loss of CENP-A peak 1), and E2 (loss of CENP-A peak 3) demonstrate lack of genetic rearrangements occurring at the site of NeoCEN4 in clones or parental population. Height of colored lines indicates what type of rearrangement they signify.

**Figure S2. Whole genome sequencing identifies loss of heterozygosity at NeoCEN4, related to Figures 1-2. A.** Density of variants between maternal and paternal haplotypes in PD-NC4 parental cells and clones reveals loss of variation at NeoCEN4, but this variation loss is not unique in the ∼ 5 Mb surrounding NeoCEN4. **B.** Expanded view of variant allele frequency in WGS (top) and CENP-A ChIP-seq (bottom) reads at NeoCEN4 in parental PD-NC4 cells and clones illustrates the loss of heterozygosity between maternal and paternal alleles at NeoCEN4 locus. **C.** Identification of SNPs at NeoCEN4 in parental PD-NC4 cells and clones, highlighting the inability to assess the epigenetics of the single copy of chromosome 4 containing NeoCEN4 by phasing short reads between alleles.

**Figure S3. DiMeLo-seq for CENP-A, DNA methylation, and H3K9me3 in PD-NC4 clones when young at NeoCEN4 and CEN4, related to Figures 3-4. A.** Single molecule DiMeLo-seq tracks illustrating probability (≥0.8) of CENP-A (red), DNA methylation (blue) and K3K9me3 (green) at individual bases in PD-NC4 clone D2 when young across NeoCEN4. **B.** Single molecule DiMeLo-seq tracks illustrating probability (≥0.8) of CENP-A (red), DNA methylation (blue) and K3K9me3 (green) at individual bases in PD-NC4 clone D2 after 12 months across CEN4. In **A-B**, aggregate tracks show mA/A and mCpG/CpG in the targeted sample in 5 kb bins. **C, D.** DiMeLo-seq for H3K9me3 enrichment in the 7 Mb surrounding NeoCEN4’s active **(C)** and inactive **(D)** locus in PD-NC4 clones. In **C-D**, Grey shadow marks NeoCEN4 locus. DiMeLo-seq tracks in **A** and **B** depict the average fraction of modified bases over 5000 bp windows with probability ≥ 0.8.

**Figure S4. Epigenetics of the inactive NeoCEN4 locus over time, related to Figures 3, 5. A,** 1. **B.** DiMeLo-seq for CENP-A (red), DNA methylation (blue) and H3K9me3 (green) at the inactive NeoCEN4 locus in PD-NC4 clones when young **(A)** and after 12 months (**B).** Purple bars in **A** and **B** highlight the location of LINE1 element refractive to acquiring CENP-A, which is not enriched for H3K9me3 in the inactive NeoCEN4 locus. **C.** DiMeLo-seq after 12 months at the inactive NeoCEN4 locus on chromosome 4 haplotype 2 in clone D2, overlaid on DiMeLo-seq data from clone D2 young. **D.** DiMeLo-seq for CENP-A (red) and DNA methylation (blue) at a subclone isolated from PD-NC4 clone C2 population after 12 months in culture confirms uncoupling of CENP-A enrichment from DNA methylation dips at NeoCEN4. DiMeLo-seq tracks in **A-D** depict the average fraction of modified bases over 5000 bp windows with probability ≥ 0.8.

**Figure S5. CEN4 locus epigenetics and evolution of NeoCEN4 in PD-NC4 clones over 12 months, related to Figures 5-6. A.** DiMeLo-seq for CENP-A (red), DNA methylation (blue) and H3K9me3 (green) at the inactive and active CEN4 locus in PD-NC4 clones C2 and E2. Horizontal grey dashed line indicates CENP-A enrichment threshold. Grey box indicates location of CENP-A enrichment and active HOR1. DiMeLo-seq tracks depict the average fraction of modified bases over 5000 bp windows with probability ≥ 0.8. **B.** CENP-A ChIP-seq or CUT&RUN for PD-NC4 parental cells and single-cell derived clones when young and after 6 and 12 months in culture. Clone tracks are colored according to their CENP-A positional phenotype assigned when young (Blue = three CENP-A peaks, Green = Loss of CENP-A peak 1, Red = Loss of CENP-A peak 3). Grey shadows represent the position of CENP-A peaks in young parental PD-NC4 cells. Purple shadows mark areas enriched for LINE1 elements in PD-NC4 cells. Black bars show location of significantly enriched CENP-A peaks (p<0.00001, fc ≥ 20). **C.** Average m6A/A (CENP-A enrichment) and variation in m6A/A among DiMeLo-seq reads aligned across active CEN4 (Haplotype 2) in PD-NC4 clone C2 at young and 12 months timepoints. Positions are binned across 100 bp. **D.** Quantification of variation in m6A/A (CENP-A enrichment) per position (100bp bins) at CEN4 in **C** in C2 Young and 12 months datasets. Lines in violin plots represent median variation values. IQR, Inter-quartile range. STD, Standard deviation.

**Figure S6. Characterization of CENP-A and HJURP overexpression in PD-NC4 CENP-A^LAP-eYFP^/HJURP^LAP-mCherry^ clones, related to Figure 7. A.** Western blot for CENP-A in PD-NC4 parental (Ctrl) cells and in CENP-A^LAP-eYFP^/HJURP^LAP-mCherry^ clones. CENP-A^LAP-eYFP^ and endogenous CENP-A levels were used to quantify total CENP-A expression in clones. Selected clones are highlighted in yellow. Graphs depict CENP-A fold change relative to parental PD-NC4 cell controls, with fold change values above bars. **B.** Immunofluorescence for HJURP in PD-NC4 CENP-A^LAP-eYFP^/HJURP^LAP-mCherry^ clones. **C.** Graph depicts fold change of HJURP mean nuclear intensity in clones relative to parental PD-NC4 (Ctrl), with fold change values above bars. **D.** Quantification of chromatin significantly enriched for CENP-A (p<0.00001, fc ≥ 20) at HOR1 of CEN4 in CENP-A/HJURP overexpressing PD-NC4 clones over 12 months. **E.** CENP-A CUT&RUN for CENP-A/HJURP overexpressing PD-NC4 clones aligned to PD-NC4 chromosome 4 assembly at 6 months timepoint at CEN4. Colored bar above tracks indicates positions of HOR1 and active centromere region at CEN4.

**Figure S7. Human centromeres and neocentromeres are epigenetically heterogeneous and exhibit differential epigenetic stability over prolonged proliferation, related to Figures 1-7.** Model depicting epigenetic heterogeneity which is a general feature of human centromeres and human neocentromeres within a cell population. For at least 30 cell cycles, heterogenous centromeres and neocentromeres are stably maintained within their epigenetic borders. Over prolonged proliferation, centromeres naturally evolve epigenetically but are maintained within their epigenetic borders. Conversely, over prolonged proliferation neocentromeres acquire increased epigenetic heterogeneity and fail to maintain their epigenetic borders. Overexpression of CENP-A/HJURP decreases CENP-A maintenance at centromeres but has a more dramatic effect at neocentromeres where it expands, destabilizes and causes relocation of the neocentromere over prolonged proliferation.

## Supplemental Table Legends

**Table S1. Percentage of PD-NC4 metaphases with NeoCEN4, related to Figure 1.** Quantification of NeoCEN4 presence in metaphases of PD-NC4 parental cell and clone populations.

**Table S2. Percentage of IMS13q metaphases with NeoCEN13, related to Figure 1.** Quantification of NeoCEN13 presence in metaphases of IMS13q parental cell and clone populations.

**Table S3.** P**ercentage of BBBq metaphases with NeoCEN13, related to Figure 1.** Quantification of NeoCEN13 presence in metaphases of BBB parental cell and clone populations.

**Table S4. PD-NC4 total neocentromere length, related to Figures 1-2.** Quantification of length of significant (fc > 20, p<0.00001) CENP-A enriched chromatin at NeoCEN4 in PD-NC4 parental cells and clones at young, 6 months, and 12 months timepoints.

**Table S5. IMS13q total neocentromere length, related to Figure 1.** Quantification of length of significant (fc > 20, p<0.00001) CENP-A enriched chromatin at NeoCEN13 in IMS13q parental cells and clones at young timepoint.

**Table S6. BBB total neocentromere length, related to Figure 1.** Quantification of length of significant (fc > 20, p<0.00001) CENP-A enriched chromatin at NeoCEN13 in BBB parental cells and clones at young timepoint.

## Methods

### Cell culture

PD-NC4 fibroblasts were previously immortalized and partially transformed by expressing hTERT and oncogenic KRASV12 (Mahlke et al., 2023) and maintained in DMEM medium (Gibco) containing 10% fetal bovine serum (Omega Scientific), 100U/ml penicillin, 100U/ml streptomycin and 2mM l-glutamine at 37°C in a 5% CO2 atmosphere with 21% oxygen. IMS13q lymphocytes and BBB fibroblasts immortalized by ectopic retroviral expression of human telomerase (*hTERT)* (Alonso et al., 2003) were a gift from Peter E. Warburton and were maintained in the same conditions, except for 15% fetal bovine serum used for IMS13q lymphocytes. Cells were maintained and split every 2-4 days according to ATCC recommendations. PD-NC4 cells overexpressing CENP-A or CENP-A and HJURP were generated via lentiviral transduction of lentiviruses expressing LAPeYFP tagged-CENP-A and/or LAPmCherry tagged-HJURP (LAP tag consists of 6x histidine, protease cleavage site and EYFP or mCherry) under a CMV promoter that were cloned into pSMPUW-IRES-NEO (VPK-216, Cell Biolabs). Single-cell derived clones were obtained by limited dilution or by sorting into 96-well cell culture plates and were first frozen down as ‘young’ 8-9 weeks (∼40 cell cycles) after limited dilution. PD-NC4 single-cell derived clones were classified into CENP-A positional phenotypes based on their overall CENP-A distribution and MACS2 peak calls (see below) when young. PD-NC4 clones were maintained in culture for an additional 12 months from the ‘Young’ time point (for a total of ∼280 cell cycles). To increase recovery of single-cell derived IMS13q lymphocytes clones after sorting, single lymphocytes were fed with lymphocyte conditioned medium (DMEM, 15% FBS, 100U/ml penicillin, 100U/ml streptomycin, 2mM l-glutamine) harvested from parental IMS13q lymphocytes after 2 days and passed through a 0.22 um filter.

### Immunofluorescence, Chromosome Spreads and IF-FISH

Immunofluorescence on chromosome spreads was done by pre-treating the cells with Karyomax (Thermo Fisher Scientific, 1:100) for 3-6 hours. Cell pellets were resuspended in DPBS and pre-warmed 37°C 75mM KCl was added dropwise to resuspended cells with gentle vortexing. Cells were cytospun onto slides at 1800 RPM for 10 minutes, then fixed with 4% formaldehyde. All other cells were fixed with 4% formaldehyde, permeabilized with 0.3% PBS-T, and blocked with triton block. Incubations with primary antibodies were conducted in triton block for 1 hr at room temperature using the following antibodies: CENP-A (Abcam ab13939, 1:1000), CENP-B (Abcam ab25734, 1:1000). NeoCEN4 was identified by non-colocalization of CENP-A and CENP-B. For analysis of NeoCEN4 fragility, cell cultures were first incubated with 0.2uM aphidicolin (Sigma-Aldrich A0781) or DMSO (Sigma-Aldrich D2650) for 24 hours, followed by metaphase spreads preparation. For IF-FISH metaphase cells were prepared as described above and then processed as described previously (Ly et al., 2019) with FISH probe to NeoMM region (Empire Genomics, RP11-752M1, hg19/hg38 Chr4:135907278-136098599 / Chr4:134986123-135177444). Images were captured at 90X magnification with 11 x 0.2 µm z-sections using a Nikon Eclipse Ti2 inverted microscope and processed with NIS Element and FIJI.

### Centromere quantification

Un-deconvolved images were prepared as maximum intensity projections and CENP-A and CENP-B intensity were measured for all centromeres in 50 randomly selected interphase cells for each PD-NC4 clone using CRaQ (Bodor et al., 2012) as described previously (Fachinetti et al., 2015). Neocentromeres in each cell were identified by presence of CENP-A and absence of CENP-B staining (<10 F.U.). Cells where neocentromeres could not be identified were discarded from the dataset. Clones were grouped according to CENP-A deposition patterns at the neocentromere for statistical analysis. Three CENP-A deposition groups were defined: 1) loss of peak 1, 2) maintaining all three peaks, and 3) loss of peak 3. CENP-A fluorescence was converted to molecules of CENP-A by normalizing it to the number of CENP-A molecules previously calculated for centromeres and neocentromere in the PD-NC4 cell line (Bodor et al., 2012).

### Micronuclei quantification

Images were deconvolved and prepared as maximum intensity projections. The number of nuclei and frequency of micronuclei were measured in three replicates of twenty images captured at 90x for each sample. Micronuclei were evaluated for the presence of CENP-A and CENP-B to determine whether they contained NeoCEN4 (non-colocalizing CENP-A and CENP-B or CENP-A only), a centromere (co-localizing CENP-A and CENP-B), or a DNA fragment without centromeric regions (no CENP-A or CENP-B). Clones were grouped according to CENP-A deposition patterns at NeoCEN4 described above for statistical analysis. The rate of micronuclei formation was calculated as the percentage of cells with micronuclei out of the total number of cells quantified for each replicate. The percentage of micronuclei containing NeoCEN4 was calculated out of the total number of micronuclei quantified. Statistical analysis and data visualization was performed using GraphPad Prism software, and one-way ANOVA tests (p=0.05) were performed to determine statistical significance.

### Sister chromatid cohesion

Distance between sister chromatids at neocentromeres was calculated on metaphase spreads by measuring distance between metaphase CENP-C foci at centromeres that lacked CENP-B (<10 F.U.). For each clone, a total of 15 neocentromeres and 75 centromeres were measured. To control for clonal effects and spread preparations unrelated to NeoCEN4 function, distance between CENP-C foci at NeoCEN4 was normalized to distance between CENP-C foci at other centromeres within the same cell spread and expressed as a ratio to normal centromere average. Clones were grouped according to the three CENP-A deposition patterns at NeoCEN4 described above for statistical analysis. One-way ANOVA tests (p=0.05) were performed to determine statistical significance.

### Cell growth

PD-NC4 parental cells and single-cell derived clones were plated in triplicate into 12-well plates at a density of 50K cells per well and allowed to grow for 14 days. Individual wells were harvested and counted every two days using a Nexcelom cellometer. One-way ANOVA tests (p=0.05) were performed to determine statistical significance.

### PD-NC4 CENP-A^LAP-eYFP^ and HJURP^LAP-mCherry^ clones and characterization

PD-NC4 cells were stably transfected with lentiviruses packaged with CENP-A^LAP-eYFP^ and HJURP^LAP-mCherry^ expression constructs, then sorted with flow cytometry to select for a double positive (eYFP+, mCherry+) population prior to single-cell cloning. Immunoblotting was performed as described previously (Nechemia-Arbely et al., 2019) using antibodies for CENP-A (ab13939, Abcam; 1:200) to quantify CENP-A overexpression level in individual clones. CENP-A expression levels were measured by quantifying expression of endogenous CENP-A and CENP-A^LAP-eYFP^. IF for HJURP (HJURP3399, 1:500, Covance, Denver PA) (Foltz et al., 2009). was used to quantify HJURP overexpression level in individual clones and performed as detailed above. Mean nuclear HJURP intensity for >50 cells was measured for each clone and parental, non-overexpressing PD-NC4 cells, with overexpression values calculated as fold change increase relative to parental HJURP levels.

### Whole genome sequencing

Genomic DNA was prepared using the Qiagen DNeasy kit (#69504) and sequencing was performed using 150 bp PE sequencing on a NovaSeq 6000 instrument. To identify whether NeoCEN4 contains any insertions/deletions in individual clones, reads from whole-genome sequencing and CENP-A ChIP-seq were aligned to the GRCh38 reference genome using bwa-mem (0.7.17) (Li and Durbin, 2009). SNPs and indels were called using VarDict (Lai et al., 2016). Structural variants were called using both brass (6.3.3) and gridss (2.12.2) structural variant callers (Cameron et al., 2021; Cameron et al., 2017; Campbell et al., 2008). Brass was run with Ensembl version 91 and a synthetic sample statistic with rho 1.0. Ploidy 2.0 GenderChr Y GenderChrFound Y. Gridss output was annotated using RepeatMasker (4.1.2) with the Dfam (3.3) library and NCBI/RMBLAST (2.11.0) engine and filtered with GRIPSS (1.11). Sample depth ratios were calculated using Cobalt (1.11) and B allele frequencies estimated with Amber (3.5) in tumor-only mode. Segmental copy numbers were estimated using Purple (3.8.4), tumor-only mode with the cobalt, amber, and gridss structural variant calls as input (Cameron et al., 2019).

### ChIP-seq

Chromatin was extracted from 1×10^8^ PD-NC4 nuclei and prepared for CENP-A ChIP-seq followed by DNA extraction as previously described (Nechemia-Arbely et al., 2019). ChIP libraries were prepared using NEB Ultra II (Cat #E7103L) following NEBNext protocols with minor modifications. To reduce biases induced by PCR amplification of a repetitive region, libraries were prepared from 80-100 ng of input or ChIP DNA. The DNA was end-repaired and A-tailed and NEBNext adaptors (Cat #E7335S) were ligated. The libraries were PCR-amplified using only 5-7 PCR cycles since the starting DNA amount was high. Libraries were run on a 2% agarose gel and size selected for 200-350 bp. Resulting libraries were sequenced using 150 bp, paired-end sequencing on a HiSeq X instrument per manufacturer’s instructions (Illumina, San Diego, CA).

Reads generated from PD-NC4 CENP-A ChIP-seq were processed with Cutadapt (v.2.10) to remove adapters and retain reads with minimum length of 20 bp, then reads were aligned to the CHM13 whole-genome assembly v2.0 (GCF_009914755.1) using BWA (v0.7.17) with the following parameters: bwa mem -t 2 -k 50 -c 1000000 [index] [read1.fastq] [read2.fastq] for paired-end data. The resulting SAM files were converted to BAM format using SAMtools (v1.9) with FLAG score 2308 to remove unmapped, secondary, and supplemental alignments. Normalized bigWigs were generated with deeptools (v.3.3.0) using the following command: bamCoverage -b sample.sort.bam -o sample.bw --scaleFactor X -p max/2, where sequencing coverage was used to calculate scaling factors for all samples. Significantly enriched CENP-A peaks were determined with MACS2 (v2.2.7.1) using default parameters, -g 3.03e9 and -q 0.00001 and subsequently filtered to remove peaks with fold change <20.

### CUT&RUN

CUT&RUN for CENP-A was performed with 1 x10^6^ cells per sample as previously described (Skene and Henikoff, 2017). Cells were incubated with CENP-A antibody (ab13939, Abcam) overnight at 4°C followed by secondary antibody for rabbit IgG (ab46540, Abcam) for 1 hour at 4°C and incubation with pA/G-MNase (Epicypher) at 4°C for 1 hour. Released chromatin fragments were prepared for sequencing using the NEB Ultra II library preparation kit (Cat #E7103L). All samples were amplified for 12 PCR cycles then run on a 2% agarose gel and size selected for mononucleosome fragments of 275-350 bp. Excised fragments were purified by gel extraction (Qiagen, #28704). Resulting libraries were sequenced using 150 bp, paired-end sequencing on a HiSeq X instrument per manufacturer’s instructions (Illumina, San Diego, CA).

Reads generated from CUT&RUN were processed with Cutadapt (v.2.10) to remove adapters and retain reads with minimum length of 20 bp, then reads were aligned to the CHM13 whole-genome assembly v2.0 (GCF_009914755.1) using bowtie2 (v2.4.1) with the following parameters: --end-to-end --very-sensitive --no-mixed --no-discordant -I 10 -X 700 --dovetail -p 8. The resulting SAM files were converted to BAM format using SAMtools (v1.9) with FLAG score 2308 to remove unmapped, secondary, and supplemental alignments. Normalized bigWigs were generated with deeptools (v.3.3.0) using the following command: bamCoverage -b sample.sort.bam -o sample.bw --scaleFactor X -p max/2, where sequencing coverage was used to calculate scaling factors for all samples. Significantly enriched CENP-A peaks were determined with MACS2 (v2.2.7.1) using default parameters, -g 3.03e9 and -q 0.00001 and subsequently filtered to remove peaks with fold change <20. Peaks were also called with SEACR in non-stringent mode using cutoff threshold 0.00001. SEACR and MACS2 peaks were then intersected with bedtools (v2.29.0) using -f 0.5 for sample sets containing a mix of ChIP-seq and CUT&RUN datasets.

### DiMeLo-seq

DiMeLo-seq was performed as previously described (Altemose et al., 2022b) for CENP-A (ADI-KAM-CC006-E, Enzo) and H3K9me3 (ab8898, Abcam), with slight modifications to the protocol to maximize fragment length. Samples were prepared from 10 x 10^6^ cells and washed and permeabilized cell pellets were resolved in 400 ul Tween-Wash containing primary antibody at 1:50. After washing, CENP-A samples were resuspended in 400 ul Tween-Wash with 200 nM mouse Hia5 nanobody (kindly provided by Nick Altemose, (Gamarra et al., 2025)) and H3K9me3 samples were resuspended in 400 ul Tween-Wash with 200 nM pA-Hia5 (kindly provided by Nick Altemose). After Hia5 activation, modified UHMW DNA was extracted using the NEB Monarch UHMW DNA Extraction Kit (insert here) with slight modifications to the protocol. Cells were resuspended in 40ul PBS then combined with 1.8 ml NEB Monarch HMW gDNA tissue lysis buffer and 60 ul proteinase K. DNA yield was quantified prior to library prep with the Qubit dsDNA BR Assay Kit (Invitrogen, Q32850). Library preparation was performed with the Nanopore Ultra-Long DNA Sequencing Kit (SQK-ULK001). Sequencing was performed with R9 flow cells on a Promethion P2 using adaptive sampling to enrich for centromeres and NeoCEN4. After 24 hours, libraries were recovered from flow cells, flow cell washes were done with the Nanopore Flow Cell Wash Kit XL (EXP-WSH004-XL), and libraries were reloaded and run for another 24 hours.

Basecalling for R9 flow cells was performed using guppy (v6.1.2_gpu) and modified bases were identified with 5mC (res_dna_r941_min_modbases-all-context_v001) and CpG (res_dna_r941_min_modbases_5mC_CpG_v001) models with a methylation threshold of 0.05. Basecalling for R10 flow cells was performed using Dorado (v0.5.3) and modified bases were identified with calls to internal hac, 5mC_5hmC, and 6mA models. Reads were aligned to an assembly of the maternal and paternal PDNC4 chromosome 4 (kindly provided by Rachel O’Neill and Savanna Hoyt, accession number SAMN46577039) using winnowmap (v2.03) for repetitive centromere regions or using minimap2 (v2.24) for the non-repetitive NeoCEN4 locus with the following parameters: -ax map-ont --cs --eqx -Y -L -p 0.1 -I8g. The resulting SAM files were converted to BAM format using SAMtools (v1.12) with FLAG score 2308 to remove unmapped, secondary, and supplemental alignments. Modified base tags were combined with winnowmap or minimap2 alignments using samtools (v1.12) and a merging script (kindly provided by Nick Altemose). Resulting alignments were filtered to isolate haplotypes using samtools (v1.12) and to remove low quality basecalls using a python script (kindly provided by Yuan Xu, Miga Lab). Visualizations were generated with the Dimelo package using probability threshold ≥ 0.8 (Maslan et al., 2024). Threshold for CENP-A enrichment (horizontal grey dashed line in Fig. 3C, D) was determined by analyzing background noise at the active CEN4 and identifying mA/A ratio levels that represent true CENP-A enrichment (Fig. 4B). Repeat annotations were generated by S. Hoyt using the repeat annotation pipeline in (Hoyt et al., 2022). Variation between CENP-A (m6A/A) position among individual DiMeLo-seq reads was calculated over 100bp bins and plotted with a custom python script available at https://github.com/mmahlke/Dimelo-seq/blob/main/Dimelo_variation_plot_complete.py.

## Data availability

ChIP-seq and CUT&RUN sequencing data aligned to the T2Tv2.0 assembly and bed tracks for significantly enriched peaks can be accessed here:

PD-NC4 cells https://genome.ucsc.edu/s/Arbelylab/PD%2DNC4_reviewers

IMS13q cells https://genome.ucsc.edu/s/Arbelylab/IMS13q_reviewers

BBB cells https://genome.ucsc.edu/s/Arbelylab/BBB_reviewers

PD-NC4 subclones https://genome.ucsc.edu/s/Arbelylab/PD%2DNC4_subclones_reviewers

All sequencing data generated in this paper is currently being submitted to SRA at PRJNA1231498

PD-NC4 chromosome 4 assembly is available at GEO accession number SAMN46577039

