## Supplemental Figures for "Evolution and instability of human centromeres are accelerated by heterochromatin boundary loss and CENP-A overexpression"

Mahlke et al., Figure S1

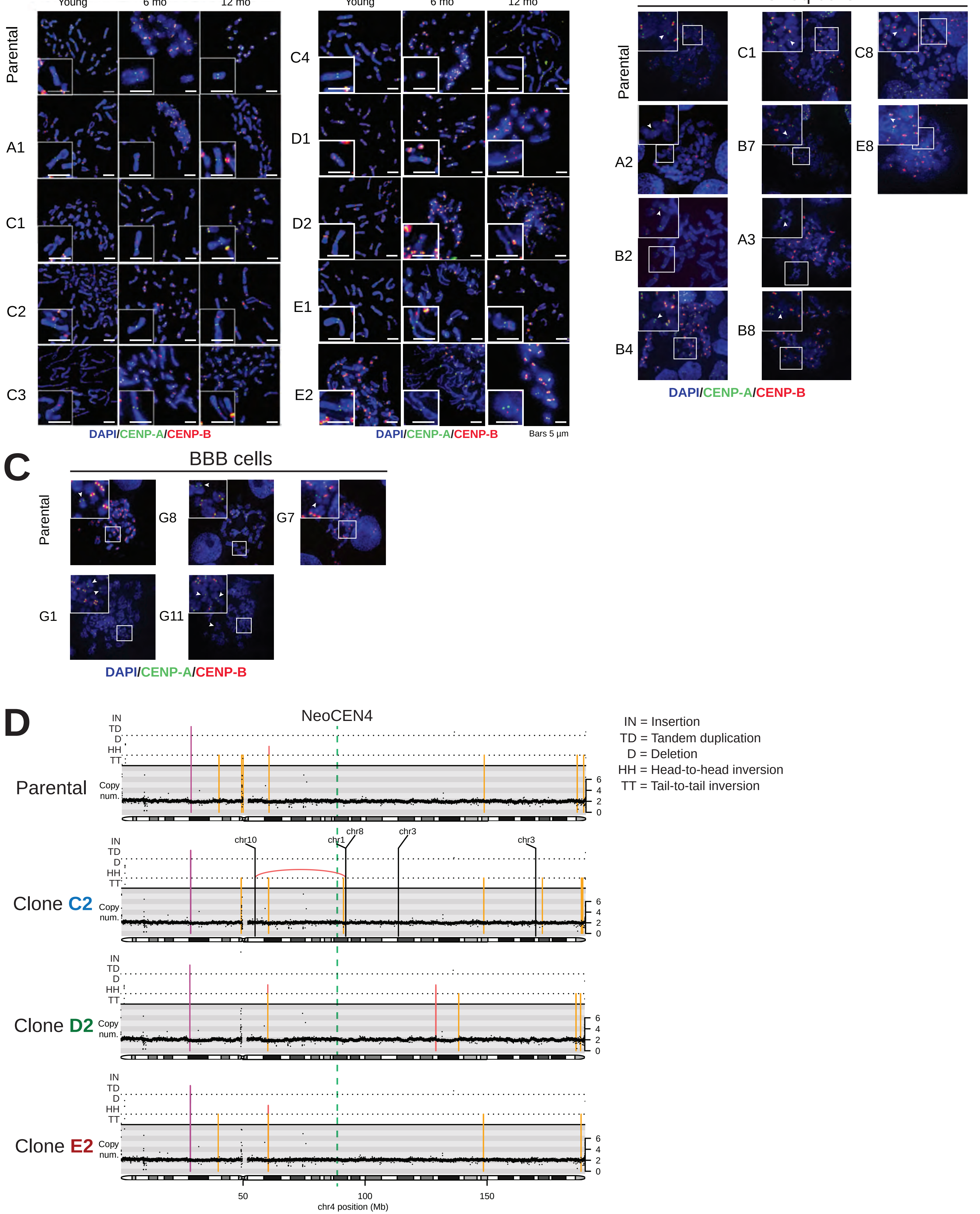


Figure S1. NeoCEN4 and NeoCEN13 are maintained within their respective populations,related to Figure 1.

Mahlke et al., Figure S2

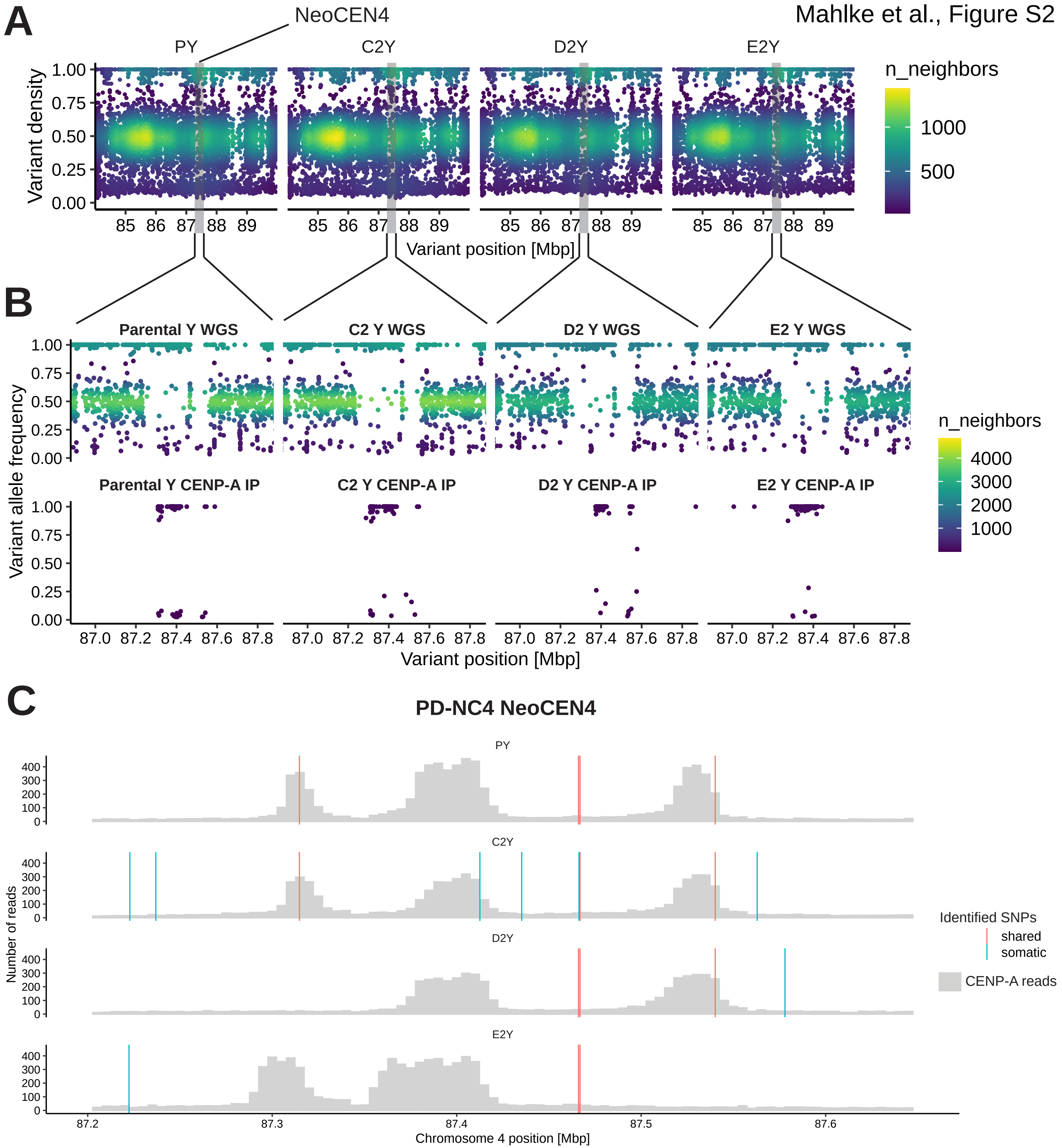


Figure S2. Whole genome sequencing identifies loss of heterozygosity at NeoCEN4, related to Figures 1-2.

Mahlke et al., Figure S3

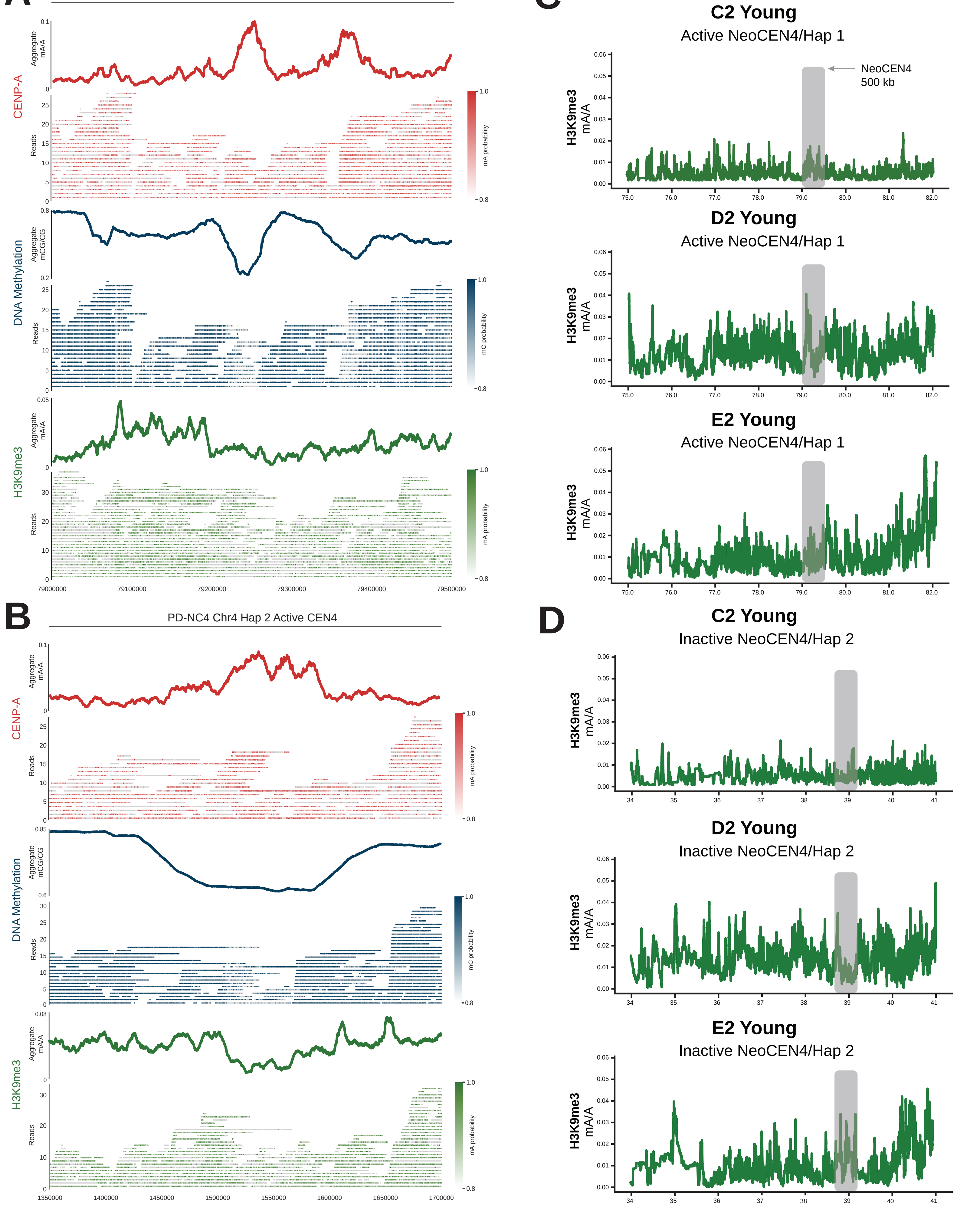


Figure S3. DiMeLo-seq for CENP-A, DNA methylation, and H3K9me3 in PD-NC4 clones when young at NeoCEN4 and CEN4, related to Figures 3-4.

Mahlke et al., Figure S4

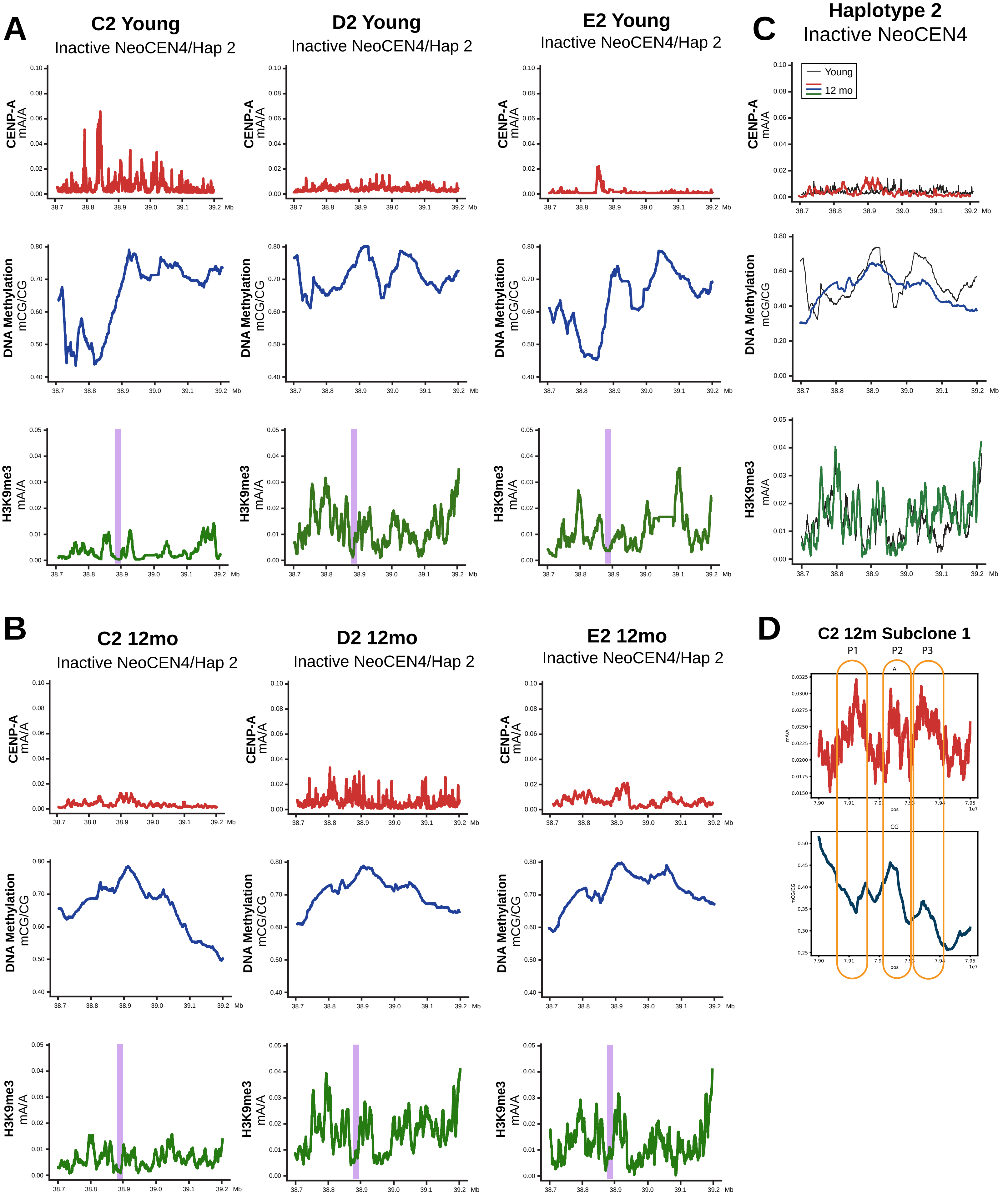


Figure S4. Epigenetics of the inactive NeoCEN4 locus and larger epigenetic footprint, related to Figures 3, 5.

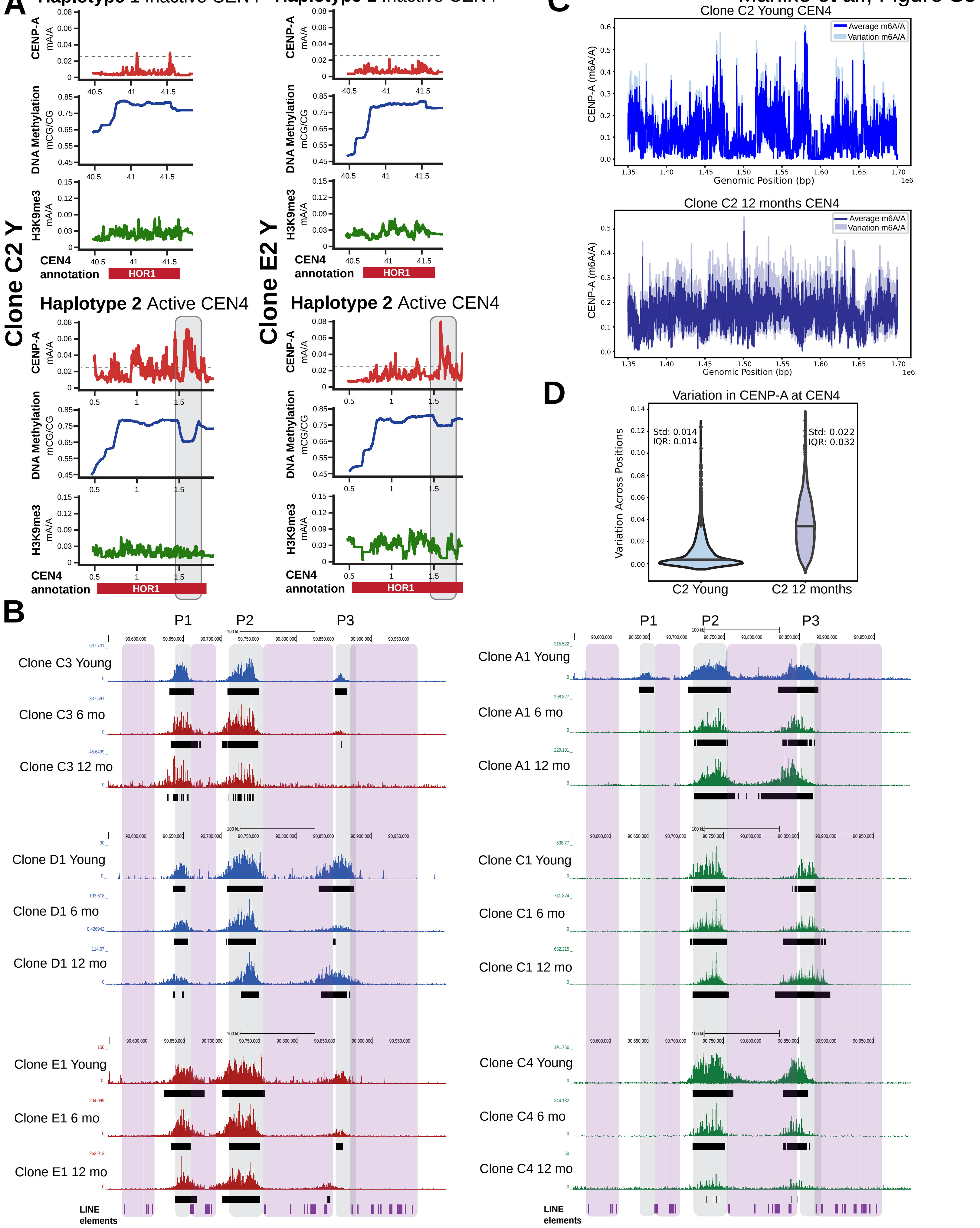


Figure S5. CEN4 locus epigenetics and evolution of NeoCEN4 in PD-NC4 clones over 12 months, related to Figures 5-6.

Mahlke et al., Figure S6

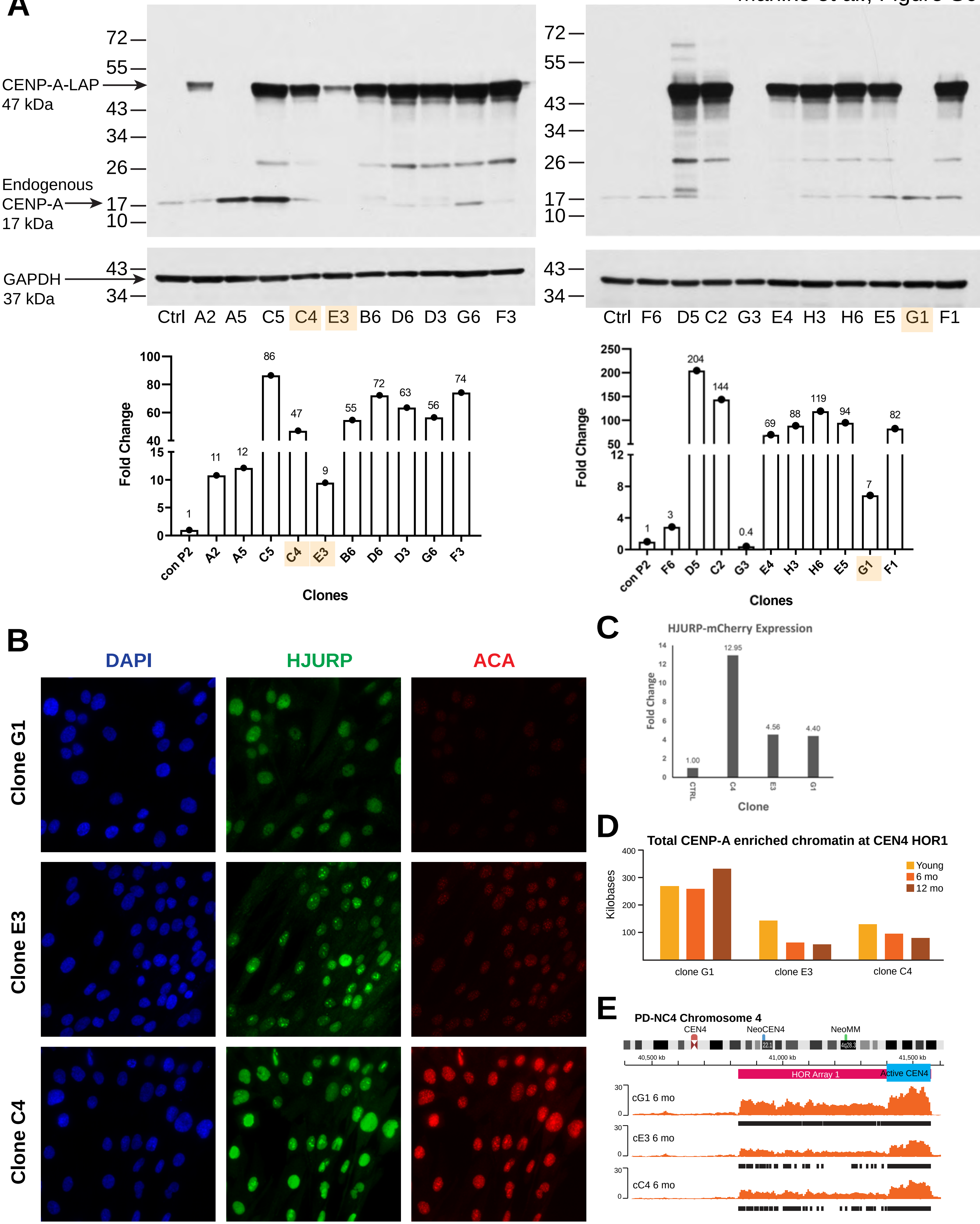


Figure S6. Characterization of CENP-A and HJURP overexpression in PD-NC4 CENP-A-LAP-eYFP/HJURP-LAP-mCherry clones, related to Figure 7.

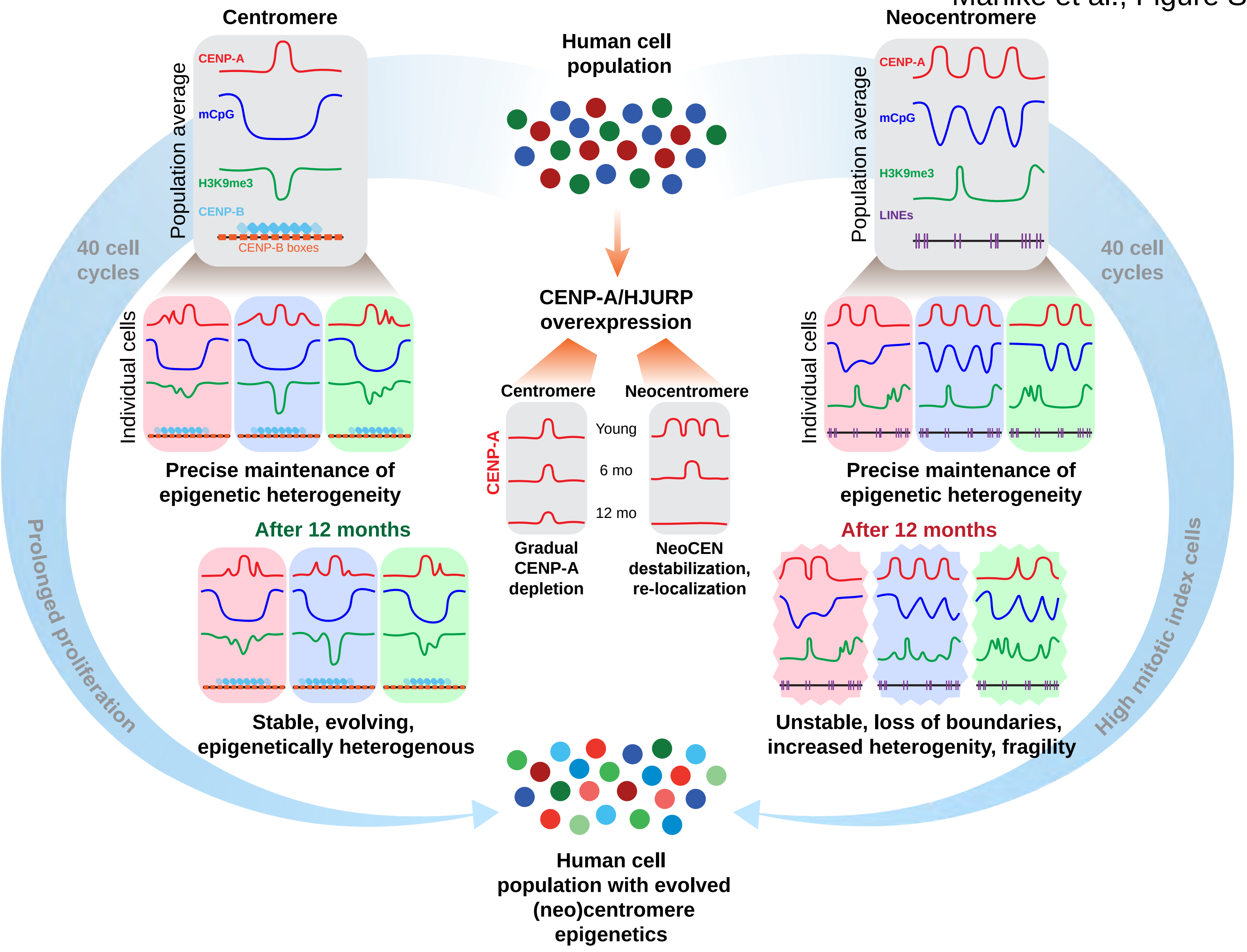


Figure S7. Human centromeres and neocentromeres are epigenetically heterogeneous and exhibit differential epigenetic stability over prolonged proliferation, related to Figures 1-7.

**Table S1. Percentage of PD-NC4 metaphases with NeoCEN4, related to Figure 1.**

Quantification of NeoCEN4 presence in metaphases of PD-NC4 parental cell and clone populations.

| Age | | Young | 6 months | 12 months |
|---|---|---|---|---|
| PD-NC4 Cell line | Parental | 100 | 100 | 60 |
| | A1 | 100 | 86 | 80 |
| | C1 | 100 | 100 | 95 |
| | C2 | 100 | 75 | 100 |
| | C3 | 100 | 86 | 90 |
| | C4 | 55 | 35 | 20 |
| | D1 | 100 | 70 | 85 |
| | D2 | 100 | 100 | 85 |
| | E1 | 90 | 100 | 90 |
| | E2 | 100 | 100 | 74 |

**Table S2. Percentage of IMS13q metaphases with NeoCEN13, related to Figure 1.**

Quantification of NeoCEN13 presence in metaphases of IMS13q parental cell and clone populations.

| Age | | Young | Selected for CUT&RUN |
|---|---|---|---|
| IMS13q Cell line | Parental | 100 | YES |
| | B4 | 95 | |
| | B7 | 90 | |
| | A4 | 100 | |
| | C1 | 95 | |
| | C8 | 95 | |
| | B8 | 90 | |
| | B2 | 80 | |
| | A3 | 85 | |
| | E8 | 80 | |
| | A8 | 60 | NO |
| | A7 | 75 | |
| | A2 | 35 | |
| | A1 | 50 | |
| | A6 | 60 | |
| | B3 | 60 | |

**Table S3. Percentage of BBBq metaphases with NeoCEN13, related to Figure 1.**

Quantification of NeoCEN13 presence in metaphases of BBB parental cell and clone populations.

| Age | | Young | Selected for CUT&RUN |
|---|---|---|---|
| BBBq Cell line | Parental | 90 | YES |
| | G1 | 75 | |
| | G7 | 95 | |
| | G8 | 90 | |
| | G11 | 100 | |
| | H2 | 60 | NO |
| | B2-3 | 30 | |
| | H11 | 25 | |
| | G2-2 | 20 | |
| | B9 | 20 | |

**Table S4. PD-NC4 total neocentromere length, related to Figures 1-2.**

Quantification of length of significant (fc > 20, p<0.00001) CENP-A enriched chromatin at NeoCEN4 in PD-NC4 cells and clones at young, 6 months, and 12 months timepoints.

| Age | Cell Line | Peak lengths (kb) | | | Total length (kb) |
|---|---|---|---|---|---|
| | | Peak 1 | Peak 2 | Peak 3 | |
| Young | Parental | 16.43 | 45 | 22.59 | 84.02 |
| | C2 | 21.82 | 44.01 | 27.61 | 93.44 |
| | D1 | 20.86 | 44.71 | 25.44 | 91.01 |
| | C3 | 30.67 | 44.42 | 18.81 | 93.9 |
| | A1 | 11.52 | 50.88 | 44.31 | 106.71 |
| | C1 | 0 | 53.11 | 49.38 | 102.49 |
| | C4 | 0 | 63.76 | 43.52 | 107.28 |
| | D2 | 0 | 45.25 | 36.37 | 81.62 |
| | E1 | 28.48 | 45.65 | 0 | 74.13 |
| | E2 | 30.35 | 61.31 | 0 | 91.66 |
| 6 months | Parental | 0 | 68.85 | 0 | 94.72 |
| | C2 | 21.17 | 41.66 | 49.95 | 112.78 |
| | D1 | 20.86 | 44.71 | 25.44 | 91.01 |
| | C3 | 37.5 | 47.85 | 0 | 85.35 |
| | A1 | 0 | 51.88 | 44.8 | 96.68 |
| | C1 | 0 | 55.21 | 64.15 | 119.36 |
| | C4 | 0 | 48.14 | 39.01 | 87.15 |
| | D2 | 0 | 39.13 | 38.79 | 77.92 |
| | E1 | 29.4 | 45.65 | 10.82 | 85.87 |
| | E2 | 74.12 | 70.12 | 0 | 144.24 |
| 12 months | Parental | 43.82 | 68.28 | 0 | 123.68 |
| | C2 | 26.5 | 42.7 | 45.64 | 114.84 |
| | D1 | 23.19 | 27.71 | 43.37 | 94.27 |
| | C3 | 27.96 | 34.06 | 0 | 62.02 |
| | A1 | 0 | 50.55 | 69.2 | 119.75 |
| | C1 | 0 | 55.46 | 72.15 | 127.61 |
| | C4 | 0 | 0 | 0 | Lost @ 12 mo |
| | D2 | 0 | 20.59 | 61.51 | 82.1 |
| | E1 | 36.2 | 67.2 | 8.94 | 112.34 |
| | E2 | 42.7 | 59.88 | 0 | 102.58 |

**Table S5. IMS13q total neocentromere length, related to Figure 1.**

Quantification of length of significant (fc > 20, p<0.00001) CENP-A enriched chromatin at NeoCEN13 in IMS13q cells and clones at young timepoint.

| Age | Cell Line | Peak lengths (kb) | | | Total length (kb) |
|---|---|---|---|---|---|
| | | Peak 1 | Peak 2 | Peak 3 | |
| Young | Parental | 18.67 | 61.7 | | 80.37 |
| | A4 | 16.8 | 34.35 | 34.45 | 85.6 |
| | B2 | 23.35 | 71.83 | | 95.18 |
| | B4 | 18.06 | 71.39 | | 89.45 |
| | C1 | 13.81 | 30.8 | 38.58 | 83.19 |
| | B7 | 0 | 68.01 | | 68.01 |
| | A3 | 18.61 | 27.65 | 0 | 46.26 |
| | B8 | 18.55 | 27.58 | 0 | 46.13 |
| | C8 | 18.59 | 38.8 | 0 | 57.39 |
| | E8 | 23.31 | 34.48 | 0 | 57.79 |

**Table S6. BBB total neocentromere length, related to Figure 1.**

Quantification of length of significant (fc > 20, p<0.00001) CENP-A enriched chromatin at NeoCEN13 in BBB cells and clones at young timepoint.

| Age | Cell Line | Total length (kb) |
|---|---|---|
| Young | Parental | 66.86 |
| | G1 | 63.68 |
| | G7 | 63.54 |
| | G8 | 71.72 |
| | G11 | 59.39 |
